# Single-trial characterization of neural rhythms: potential and challenges

**DOI:** 10.1101/356089

**Authors:** Julian Q. Kosciessa, Thomas H. Grandy, Douglas D. Garrett, Markus Werkle-Bergner

## Abstract

The average power of rhythmic neural responses as captured by MEG/EEG/LFP recordings is a prevalent index of human brain function. Increasing evidence questions the utility of trial-/group averaged power estimates however, as seemingly sustained activity patterns may be brought about by time-varying transient signals in each single trial. Hence, it is crucial to accurately describe the duration and power of rhythmic and arrhythmic neural responses on the single trial-level. However, it is less clear how well this can be achieved in empirical MEG/EEG/LFP recordings. Here, we extend an existing rhythm detection algorithm (**e**xtended **B**etter **OSC**illation detection: “eBOSC”; cf. Whitten et al., 2011) to systematically investigate boundary conditions for estimating neural rhythms at the single-trial level. Using simulations as well as resting and task-based EEG recordings from a micro-longitudinal assessment, we show that alpha rhythms can be successfully captured in single trials with high specificity, but that the quality of single-trial estimates varies greatly between subjects. Despite those signal-to-noise-based limitations, we highlight the utility and potential of rhythm detection with multiple proof-of-concept examples, and discuss implications for single-trial analyses of neural rhythms in electrophysiological recordings. Using an applied example of working memory retention, rhythm detection indicated load-related increases in the duration of frontal theta and posterior alpha rhythms, in addition to a frequency decrease of frontal theta rhythms that was observed exclusively through amplification of rhythmic amplitudes.

**Highlights:**

- Traditional narrow-band rhythm metrics conflate the power and duration of rhythmic and arrhythmic periods. We extend a state-of-the-art rhythm detection method (eBOSC) to derive rhythmic episodes in single trials that can disambiguate rhythmic and arrhythmic periods.
- Simulations indicate that this can be done with high specificity given sufficient rhythmic power, but with strongly impaired sensitivity when rhythmic SNR is low. Empirically, surface EEG recordings exhibit stable inter-individual differences in α-rhythmicity in ranges where simulations suggest a gradual bias, leading to high collinearity between narrow-band and rhythm-specific estimates.
- Beyond these limitations, we highlight multiple empirical benefits of characterizing rhythmic episodes in single trials, such as (a) a principled separation of rhythmic and arrhythmic content, (b) an amplification of rhythmic amplitudes, and (c) a specific characterization of sustained and transient events.
- In an exemplary application, rhythm-specific estimates increase sensitivity to working memory load effects, in addition to indicating a frequency modulation of frontal theta rhythms through the amplification of rhythmic power.

## 1.1 Towards a single-trial characterization of neural rhythms

Episodes of rhythmic neural activity in electrophysiological recordings are of prime interest for research on neural representations and computations across multiple scales of measurement (e.g. Buzsáki, 2006; Wang, 2010). At the macroscopic level, the study of rhythmic neural signals has a long heritage, dating back to Hans Berger’s classic investigations into the Alpha rhythm (Berger, 1938). Since then, advances in recording and processing techniques have facilitated large-scale spectral analysis schemes (e.g. Gross, 2014) that were not available to the pioneers of electrophysiological research, who often depended on the manual analysis of single time series to indicate the presence and magnitude of rhythmic events. Interestingly, improvements in analytic methods still do not capture all of the information that can be extracted by manual inspection. For example, current analysis techniques are largely naïve to the specific temporal presence of rhythms in the continuous recordings, as they often employ windowing of condition- or group-based averages to extract putative rhythm-related characteristics (Cohen, 2014). However, the underlying assumption of stationary, sustained rhythms within the temporal window of interest might not consistently be met (Jones, 2016; Stokes & Spaak, 2016), thus challenging the appropriateness of the averaging model (i.e., the ergodicity assumption (Molenaar & Campbell, 2009)). Furthermore, in certain situations, single-trial characterizations become necessary to derive unbiased individual estimates of neural rhythms (Cohen, 2017). For example, this issue becomes important when asking whether rhythms appear in transient or in sustained form (van Ede, Quinn, Woolrich, & Nobre, 2018), or when only single-shot acquisitions are feasible (i.e., resting state or sleep recordings).

## 1.2 Duration as a powerful index of rhythmicity

The presence of rhythmicity is a necessary prerequisite for the accurate interpretation of measures of amplitude, power, and phase (Aru et al., 2015; Jones, 2016; Muthukumaraswamy & Singh, 2011). This is exemplified by the bias that arrhythmic periods exert on rhythmic power estimates. Most current time-frequency decomposition methods of neurophysiological signals (such as the electroencephalogram (EEG)) are based on the Fourier transform (Gross, 2014). Following Parceval’s theorem (e.g. Hansen, 2014), the Fast Fourier Transform (FFT) decomposes an arbitrary time series into a sum of sinusoids at different frequencies. Importantly, FFT-derived power estimates do not differentiate between high-amplitude transients and low-amplitude sustained signals. In the case of FFT power, this is a direct result of the violated assumption of stationarity in the presence of a transient signal. Short-time FFT and wavelet techniques alleviate (but do not eliminate) this problem by analyzing shorter epochs, during which stationarity is more likely to be obtained. However, whenever spectral power is averaged across these episodes, both high-amplitude rhythmic and low-amplitude arrhythmic signal components may once again become intermixed. In the presence of arrhythmic content (often referred to as the “signal background,” or “noise”), this results in a reduced amplitude estimate of the underlying rhythm, the extent of which relates to the duration of the rhythmic episode relative to the length of the analyzed segment (which we will refer to as ‘abundance’) (see Figure 1A). Therefore, integration across epochs that contain a mixture of rhythmic and arrhythmic signals results in an inherent ambiguity between the strength of the rhythmic activity (as indexed by power/amplitude) and its duration (as indexed by the abundance of the rhythmic episode within the segment) (see Figure 2B).

**Figure 1:**
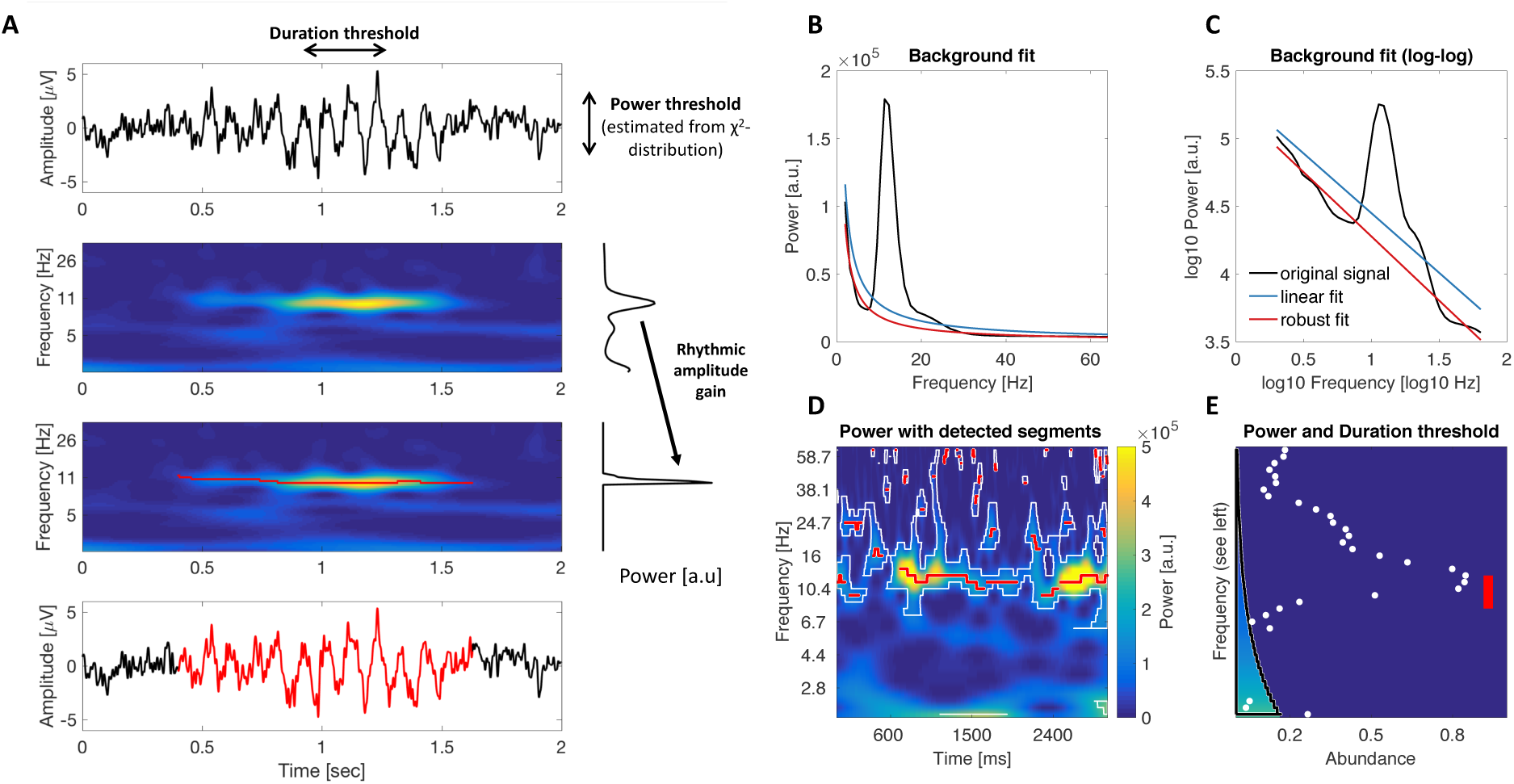
Schematic illustration of rhythm detection. (A) Average amplitude estimates (right) increase with the focus on rhythmic episodes within the averaged time interval. The left plots show simulated time series and the corresponding time-frequency power. Superimposed red traces indicate rhythmic time points. The upper right plot shows the average power spectrum averaged across the entire epoch, the lower plot presents amplitudes averaged exclusively across rhythmic time points. An amplitude gain is observed due to the exclusion of arrhythmic low amplitude time points. (B-E) Comparison of standard and extended BOSC. (B+C) Rhythms were detected based on a power threshold estimated from the arrhythmic background spectrum. Standard BOSC applies a linear fit in log-log space to define the background power, which may overestimate the background at the frequencies of interest in the case of data with large rhythmic peaks. Robust regression following peak removal alleviates this problem. (D) Example of episode detection. White borders circumfuse time frequency points, at which standard BOSC indicated rhythmic content. Red traces represent the continuous rhythmic episodes that result from the extended post-processing. (E) Applied thresholds and detected rhythmic abundance. The black border denotes the duration tshreshold at each frequency (corresponding to D), i.e., for how long the power threshold needed to be exceeded to count as a rhythmic period. Note that this threshold can be set to zero for a post-hoc characterization of the duration of episodes (see Methods 2.12). The color scaling within the demarcated area indicates the power threshold at each frequency. Abundance corresponds to the relative length of the segment on the same time scale as presented in D. White dots correspond to the standard BOSC measure of rhythmic abundance at each frequency (termed Pepisode). Red lines indicate the abundance measure used here, which is defined as the proportion of sample points at which a rhythmic episode between 8-15 Hz was indicated (shown as red traces in D).

**Figure 2:**
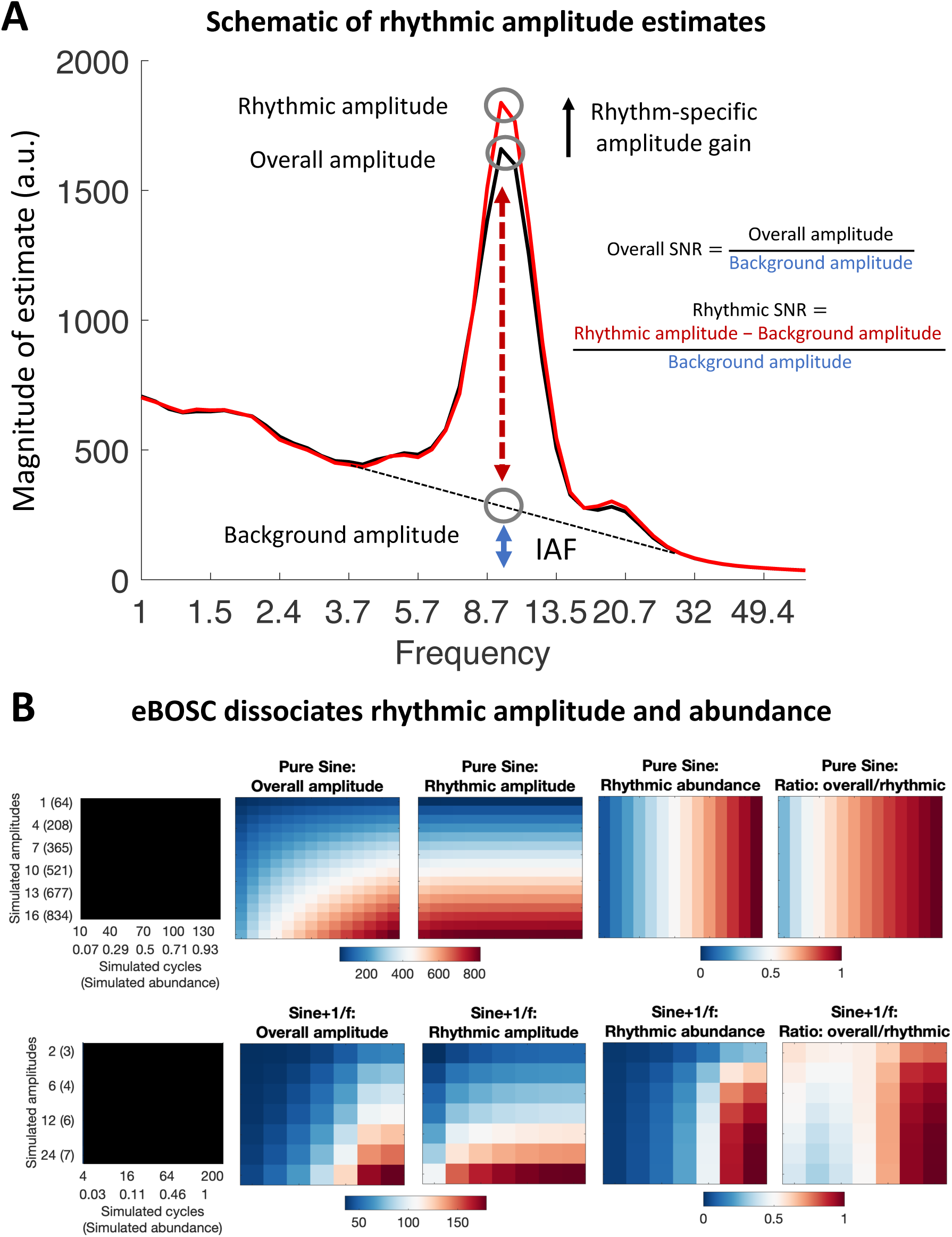
eBOSC disambiguates the magnitude and duration of rhythmic episodes. (**A**) Schema of different amplitude metrics. (**B**) Rhythm-detection disambiguates rhythmic amplitude and duration. Overall amplitudes represent a mixture of rhythmic power and duration. In the absence of noise (upper row), eBOSC perfectly orthogonalizes rhythmic amplitude from abundance. Superimposed noise leads to an imperfect separation of the two metrics (lower row). The duration of rhythmicity is similarly indicated by abundance and the overlap between rhythmic and overall amplitudes. This can be seen by comparing the two rightmost plots in each row.

Crucially, the strength and duration of rhythmic activity theoretically differ in their neurophysiological interpretation. Rhythmic power most readily indexes the magnitude of synchronized changes in membrane potentials within a network (Buzsáki, Anastassiou, & Koch, 2012), and is thus related to the size of the participating neural population. The duration of a rhythmic episode, by contrast, tracks how long population synchrony is upheld. Notably, measures of rhythm duration have recently gained interest as they may provide additional information regarding the biophysical mechanisms that give rise to the recorded signals (Peterson & Voytek, 2017; Sherman et al., 2016), for example, by differentiating between transient and sustained rhythmic events (van Ede et al., 2018).

## 1.3. Single-trial rhythm detection as a methodological challenge

In general, the accurate estimation of process parameters depends on a sufficiently strong signal in the neurophysiological recordings under investigation. Especially for scalp-level M/EEG recordings it remains elusive whether neural rhythms are sufficiently strong to be clearly detected in single trials. Here, a large neural population has to be synchronously active to give rise to potentials that are visible at the scalp surface. This problem intensifies further by signal attenuation through the skull (in the case of EEG) and the superposition of signals from diverse sources of no interest both in- and outside the brain (Schomer & Lopes da Silva, 2017). In sum, these considerations lead to the proposal that the signal-to-noise ratio (SNR), here operationally defined as the ratio of rhythmic to arrhythmic variance, may fundamentally constrain the accurate characterization of single-trial rhythms.

Following those considerations, we set out to answer the following hypotheses and questions: (1) A precise differentiation between rhythmic and arrhythmic timepoints can disambiguate the strength and the duration of rhythmicity. (2) To what extent does the single-trial rhythm representation in empirical data allow for an accurate estimation of rhythmic strength and duration in the face of variations in the signal-to-noise ratio of rhythmicity? (3) What are the empirical benefits of separating rhythmic (and arrhythmic) duration and power?

Recently, the Better OSCillation Detection (BOSC; Caplan, Madsen, Raghavachari, & Kahana, 2001; Whitten, Hughes, Dickson, & Caplan, 2011) method has been proposed to identify rhythmicity at the single-trial level. BOSC defines rhythmicity based on the presence of a spectral peak that is superimposed on an arrhythmic 1/f background and that remains present for a minimum number of cycles. Here, we extend the BOSC method (i.e., extended BOSC; eBOSC) to derive rhythmic temporal episodes that can be used to further characterize rhythmicity. Using simulations, we derive rhythm detection benchmarks and probe the boundary conditions for unbiased rhythm indices. Furthermore, we apply the eBOSC algorithm to resting- and task-state data from a micro-longitudinal dataset to systematically investigate the feasibility to derive reliable and valid indices of neural rhythmicity from single-trial scalp EEG data and to probe their modulation by working memory load.

We focus on alpha rhythms (∼8-15 Hz; defined here based on individual FFT-peaks) due to (a) their high amplitude in human EEG recordings, (b) the previous focus on the alpha band in the rhythm detection literature (Caplan, Bottomley, Kang, & Dixon, 2015; Fransen et al., 2015; Whitten et al., 2011), and (c) their importance for human cognition (Grandy, Werkle-Bergner, Chicherio, Lövdén, et al., 2013a; Klimesch, 2012; Sadaghiani & Kleinschmidt, 2016). We present examples beyond the alpha range to highlight the ability to apply eBOSC in multiple, diverse frequency ranges.

## 2. Methods

### 2.1 Study design

Resting state and task data were collected in the context of a larger assessment, consisting of eight sessions in which an adapted Sternberg short-term memory task (Sternberg, 1966) and three additional cognitive tasks were repeatedly administered. Resting state data are from the first session, task data are from sessions one, seven and eight, during which EEG data were acquired. Sessions one through seven were completed on consecutive days (excluding Sundays) with session seven completed seven days after session one by all but one participant (eight days due to a two-day break). Session eight was conducted approximately one week after session seven (M = 7.3 days, SD = 1.4) to estimate the stability of the behavioral practice effects. The reported EEG sessions lasted approximately three and a half to four hours, including approximately one and a half hours of EEG preparation. For further details on the study protocol and results of the behavioural tasks see (Grandy, Lindenberger, & Werkle-Bergner, 2017).

### 2.2 Participants

The sample contained 32 young adults (mean age = 23.3 years, SD = 2.0, range 19.6 to 26.8 years; 17 women; 28 university students) recruited from the participant database of the Max Planck Institute for Human Development, Berlin, Germany (MPIB). Participants were right-handed, as assessed with a modified version of the Edinburgh Handedness Inventory (Oldfield, 1971), and had normal or corrected-to-normal vision, as assessed with the Freiburg Visual Acuity test (Bach, 1996; 2007). Participants reported to be in good health with no known history of neurological or psychiatric incidences and were paid for their participation (8.08 € per hour, 25.00 € for completing the study within 16 days, and a performance-dependent bonus of 28.00 €; see below). All participants gave written informed consent according to the institutional guidelines of the ethics committee of the MPIB, which approved the study.

### 2.3 Procedure

Participants were seated at a distance of 80 cm in front of a 60 Hz LCD monitor in an acoustically and electrically shielded chamber. A resting state assessment was conducted prior to the initial performance of the adapted Sternberg task. Two resting state periods were used: the first encompassed a duration of two minutes of continuous eyes open (EO1) and eyes closed (EC1) periods, respectively; the second resting state was comprised of two 80 second runs, totalling 16 repetitions of 5 seconds interleaved eyes open (EO2) – eyes closed (EC2) periods. An auditory beep indicated to the subjects when to open and close their eyes.

Following the resting assessments, participants performed an adapted version of the Sternberg task. Digits were presented in white on a black background and subtended ∼2.5° of visual angle in the vertical and ∼1.8° of visual angle in the horizontal direction. Stimulus presentation and recording of behavioral responses were controlled with E-Prime 2.0 (Psychology Software Tools, Inc., Pittsburgh, PA, USA). The task design followed the original report (Sternberg, 1966). Participants started each trial by pressing the left and right response key with their respective index fingers to ensure correct finger placement and to enable fast responding. An instruction to blink was given, followed by the sequential presentation of 2, 4 or 6 digits from zero to nine. On each trial, the memory set size (i.e., load) varied randomly between trials, and participants were not informed about the upcoming condition. Also, the single digits constituting a given memory set were randomly selected in each trial. Each stimulus was presented for 200 ms, followed by a fixed 1000 ms blank inter-stimulus interval (ISI). The offset of the last stimulus coincided with the onset of a 3000 ms blank retention interval, which concluded with the presentation of a probe item that was either contained in the presented stimulus set (*positive probe*) or not (*negative probe*). Probe presentation lasted 200 ms, followed by a blank screen for 2000 ms, during which the participant’s response was recorded. A beep tone indicated the end of the trial. The task lasted about 50 minutes.

For each combination of load x probe type, 31 trials were conducted, cumulating in 186 trials per session. Combinations were randomly distributed across four blocks (block one: 48 trials; blocks two through four: 46 trials). Summary feedback of the overall mean RT and accuracy within the current session was shown at the end of each block. At the beginning of session one, 24 practice trials were conducted to familiarize participants with the varying set sizes and probe types. To sustain high motivation throughout the study, participants were paid a 28 € bonus if their current session’s mean RT was faster or equal to the overall mean RT during the preceding session, while sustaining accuracy above 90%. Only correct trials were included in the analyses.

### 2.4 EEG recordings and pre-processing

EEG was continuously recorded from 64 Ag/AgCl electrodes using BrainAmp amplifiers (Brain Products GmbH, Gilching, Germany). Sixty scalp electrodes were arranged within an elastic cap (EASYCAP GmbH, Herrsching, Germany) according to the 10% system (cf. Oostenveld, Fries, Maris, & Schoffelen, 2011) with the ground placed at AFz. To monitor eye movements, two electrodes were placed on the outer canthi (horizontal EOG) and one electrode below the left eye (vertical EOG). During recording, all electrodes were referenced to the right mastoid electrode, while the left mastoid electrode was recorded as an additional channel. Prior to recording, electrode impedances were retained below 5 kΩ. Online, signals were recorded with an analog pass-band of 0.1 to 250 Hz and digitized at a sampling rate of 1 kHz.

Preprocessing and analysis of EEG data were conducted with the FieldTrip toolbox (Oostenveld et al., 2011) and using custom-written MATLAB (The MathWorks Inc., Natick, MA, USA) code. Offline, EEG data were filtered using a 4^th^ order Butterworth filter with a pass-band of 0.5 to 100 Hz, and were linearly detrended. Resting data with interleaved eye closure were epoched relative to the auditory cue to open and close the eyes. An epoch of -2 s to +3 s relative to on- and offsets was chosen to include padding for the analysis. During the eBOSC procedure, three seconds of signal were removed from both edges (see below), resulting in an effective epoch of 4 s duration that excludes evoked components following the cue onset. Continuous eyes open/closed recordings were segmented to the cue on- and offset. For the interleaved data, the first and last trial for each condition were removed, resulting in an effective trial number of 14 trials per condition. For the task data, we analyzed two intervals: an extended interval to assess the overall dynamics of detected rhythmicity and a shorter interval that focused on the retention period. Unless otherwise noted, we refer to the extended interval when presenting task data. For the extended segments, task data were segmented to 21 s epochs ranging from -9 s to +12 s with regard to the onset of the 3 s retention interval for analyses including peri-retention data. For analyses including only the retention phase, data were segmented to -2 s to +3 s around the retention interval. Note that for all analyses, 3 s of signal were removed on each side of the signal during eBOSC detection, effectively removing the evoked cue activity (2 s to account for edge artifacts following wavelet-transformation and 1 s to account for eBOSC’s duration threshold, see section 2.6), except during the extended task interval. Hence, detected segments were restricted to occur from 1s after period onset until period offset, thereby excluding evoked signals. Blink, movement and heart-beat artifacts were identified using Independent Component Analysis (ICA; Bell & Sejnowski, 1995) and removed from the signal. Subsequently, data were downsampled to 250 Hz and all channels were re-referenced to mathematically averaged mastoids. Artifact-contaminated channels (determined across epochs) were automatically detected (a) using the FASTER algorithm (Nolan, Whelan, & Reilly, 2010) and (b) by detecting outliers exceeding three standard deviations of the kurtosis of the distribution of power values in each epoch within low (0.2-2 Hz) or high (30-100 Hz) frequency bands, respectively. Rejected channels were interpolated using spherical splines (Perrin, Pernier, Bertrand, & Echallier, 1989). Subsequently, noisy epochs were likewise excluded based on FASTER and recursive outlier detection, resulting in the rejection of approximately 13% of trials. To prevent trial rejection due to artifacts outside the signal of interest, artifact detection was restricted to epochs that included 2.4 s of additional signal around the on- and offset of the retention interval, corresponding to the longest effective segment that was used in the analyses. A further 2.65% of incorrectly answered trials from the task were subsequently excluded.

### 2.5 Rhythm-detection using extended BOSC

We applied an extended version of the Better OSCillation detection method (eBOSC; cf. Caplan et al., 2001; Whitten et al., 2011) to automatically separate rhythmic from arrhythmic episodes. The BOSC method reliably identifies rhythms using data-driven thresholds based on theoretical assumptions of the signal characteristics. Briefly, the method defines rhythms as time points during which wavelet-derived power at a particular frequency exceeds a *power threshold* based on an estimate of the arrhythmic signal background. The theoretical *duration threshold* defines a minimum duration of cycles this power threshold has to be exceeded to exclude high amplitude transients. Previous applications of the BOSC method focused on the analysis of resting-state data or long data epochs, where reliable detection has been established regardless of specific parameter setups (Caplan et al., 2001; 2015; Whitten et al., 2011). We introduce the following adaptations here (for details see section 2.6, Figure 1 & Figure S1): (1) we remove the spectral alpha peak and use robust regression to establish power thresholds; (2) we combine detected time points into continuous rhythmic episodes and (3) we reduce the impact of wavelet convolution on abundance estimates. We benchmarked the algorithm and compared it to standard BOSC using simulations (see section 2.8).

### 2.6 Specifics of rhythm-detection using extended BOSC

Rhythmic events were detected within subjects for each channel and condition. Time-frequency transformation of single trials was performed using 6-cycle Morlet wavelets (Grossmann & Morlet, 1985) with 49 logarithmically-spaced center frequencies ranging from 1 to 64 Hz. Following the wavelet transform, 2 s were removed at each segment’s borders to exclude edge artefacts. To estimate the background spectrum, the time-frequency spectra from all trials were temporally concatenated within condition and channel and log-transformed, followed by temporal averaging. For eyes-closed and eyes-open resting states, both continuous and interleaved exemplars were included in the background estimation for the respective conditions. The resulting power spectrum was fit linearly in log(frequency)-log(power) coordinates using a robust regression, with the underlying assumption that the EEG background spectrum is characterized by colored noise of the form A*f^(−α) (Buzsáki & Mizuseki, 2014; He, Zempel, Snyder, & Raichle, 2010; Linkenkaer-Hansen, Nikouline, Palva, & Ilmoniemi, 2001). A robust regression with bisquare weighting (e.g. Holland & Welsch, 2007) was chosen to improve the linear fit of the background spectrum (cf. Haller et al., 2018), which was characterized by frequency peaks in the alpha range for almost all subjects (Figure S4). In contrast to ordinary least squares regression, robust regression iteratively down-weights outliers (in this case spectral peaks) from the linear background fit. To improve the definition of rhythmic power estimates as outliers during the robust regression, power estimates within the wavelet pass-band around the individual alpha peak frequency were removed prior to fitting^1^. The passband of the wavelet (e.g. Linkenkaer-Hansen et al., 2001) was calculated as

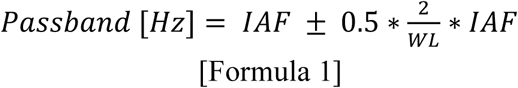

in which IAF denotes the individual alpha peak frequency and WL refers to wavelet length (here, six cycles in the main analysis). IAF was determined based on the peak magnitude within the 8-15 Hz average spectrum for each channel and condition (Grandy, Werkle-Bergner, Chicherio, Schmiedek, et al., 2013b). This ensures that the maximum spectral deflection is removed across subjects, even in cases where no or multiple peaks are present^2^. The procedure effectively removes a bias of the prevalent alpha peak on the arrhythmic background estimate (see Figure 1B and C & Figure 3C). The power threshold for rhythmicity at each frequency was set at the 95^th^ percentile of a χ^2^(2)-distribution of power values, centered on the linearly fitted estimate of background power at the respective frequency (for details see Whitten et al., 2011). This essentially implements a significance test of single-trial power against arrhythmic background power. A three-cycle threshold was used as the duration threshold to exclude transients, unless indicated otherwise (see section 2.12). The conjunctive power and duration criteria produce a binary matrix of ‘detected’ rhythmicity for each time-frequency point (see Figure S1C). To account for the duration criterion, 1000 ms were discarded from each edge of this ‘detected’ matrix.

The original BOSC algorithm was further extended to define rhythmic events as continuous temporal episodes that allow for an event-wise assessment of rhythm characteristics (e.g. duration). The following steps were applied to the binary matrix of ‘detected’ single-trial rhythmicity to derive such sparse and continuous episodes. First, to account for the spectral extension of the wavelet, we selected time-frequency points with maximal power within the wavelet’s spectral smoothing range (i.e. the pass-band of the wavelet; 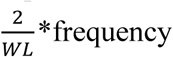; see Formula 1). That is, at each time point, we selected the frequency with the highest indicated rhythmicity within each frequency’s pass-band. This served to exclude super-threshold timepoints that may be accounted for by spectral smoothing of a rhythm at an adjacent frequency. Note that this effectively creates a new frequency resolution for the resulting rhythmic episodes, thus requiring sufficient spectral resolution (defined by the wavelet’s pass-band) to differentiate simultaneous rhythms occurring at close frequencies. Finally, continuous rhythmic episodes were formed by temporally connecting extracted time points, while allowing for moment-to-moment frequency transitions (i.e. within-episode frequency non-stationarities; Atallah & Scanziani, 2009) (for a single-trial illustration see Figures 1D and Figure S1D).

In addition to the spectral extension of the wavelet, the choice of wavelet parameter also affects the extent of temporal smoothing, which may bias rhythmic duration estimates. To decrease such temporal bias, we compared observed rhythmic amplitudes at each time point within each rhythmic episode with those expected by smoothing adjacent amplitudes using the wavelet (Figure S1E). By retaining only those time points where amplitudes exceeded the smoothing-based expectations, we removed supra-threshold time points that can be explained by temporal smoothing of nearby rhythms (e.g., ‘ramping’ up and down signals). In more detail, we simulated the positive cycle of a sine wave at each frequency, zero-shouldered each edge and performed (6-cycle) wavelet convolution. The resulting amplitude estimates at the zero-padded time points reflect the temporal smoothing bias of the wavelet on adjacent arrhythmic time points. This bias is maximal (*BiasMax*) at the time point immediately adjacent to the rhythmic on-/offset and decreases with temporal distance to the rhythm. Within each rhythmic episode, the ‘convolution bias’ of a time-frequency (TF) point’s amplitude on surrounding points was estimated by scaling the points’ amplitude by the modelled temporal smoothing bias.

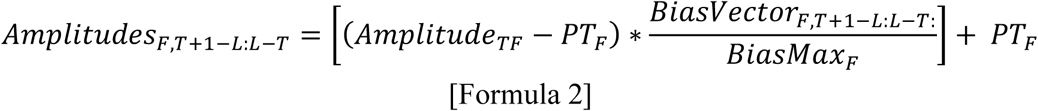

Subscripts F and T denote frequency and time within each episode, respectively. *BiasVector* is a vector with the length of the current episode (L) that is centered around the current TF-point. It contains the wavelet’s symmetric convolution bias around *BiasMax*. Note that both BiasVector and BiasMax respect the possible frequency variations within an episode (i.e., they reflect the differences in convolution bias between frequencies). The estimated wavelet bias was then scaled to the amplitude of the rhythmic signal at the current TF-point. PT refers to the condition- and frequency-specific power threshold applied during rhythm detection. We subtracted the power threshold to remove arrhythmic contributions. This effectively sensitizes the algorithm to near-threshold values, rendering them more likely to be excluded. Finally, time points with lower amplitudes than expected by the convolution model were removed and new rhythmic episodes were created (Figure S1F). The resulting episodes were again checked for adhering to the duration threshold.

As an alternative to the temporal wavelet correction based on the wavelet’s simulated maximum bias (‘MaxBias’; as described above), we investigated the feasibility of using the wavelet’s full-width half maximum (‘FWHM’) as a criterion. Within each continuous episode and for each “rhythmic” sample point, 6-cycle wavelets at the frequency of the neighbouring points were created and scaled to the point’s amplitude. We then used the amplitude of these wavelets at the FWHM as a threshold for rhythmic amplitudes. That is, points within a rhythmic episodes that had amplitudes below those of the scaled wavelets were defined as arrhythmic. The resulting continuous episodes were again required to pass the duration threshold. As the FWHM approach indicated decreased specificity of rhythm detection in the simulations (Figure S2) we used the ‘MaxBias’ method for our analyses.

Furthermore, we considered a variant where total amplitude values were used (vs. supra-threshold amplitudes) as the basis for the temporal wavelet correction. Our results suggest that using supra-threshold power values leads to a more specific detection at the cost of sensitivity (Figure S2). Crucially, this eliminated false alarms and abundance overestimation, thus rendering the method highly specific to the occurrence of rhythmicity. As we regard this as a beneficial feature, we used supra-threshold amplitudes as the basis for the temporal wavelet correction throughout the manuscript.

### 2.7 Definition of abundance, rhythmic probability and amplitude metrics

A central goal of rhythm detection is to disambiguate rhythmic power and duration (Figure 2). For this purpose, eBOSC provides multiple indices. We describe the different indices for the example case of alpha rhythms. Please note that eBOSC can be applied in a similar fashion to any other frequency range. The ***abundance*** of alpha rhythms denotes the duration of rhythmic episodes with a mean frequency in the alpha range (8 to 15 Hz), relative to the duration of the analyzed segment. This frequency range was motivated by clear peaks within this range in individual resting state spectra (Figure S4). Note that abundance is closely related to standard BOSC’s Pepisode metric (Whitten et al., 2011), with the difference that abundance refers to the duration of the continuous rhythmic episodes and not the ‘raw’ detected rhythmicity of BOSC (cf. Figure S1C and D). We further define ***rhythmic probability*** as the *across trials* probability to observe a detected rhythmic episode within the alpha frequency range at a given point in time. It is therefore the within-time, across-trial equivalent of abundance.

As a result of rhythm detection, the magnitude of spectral events can be described using multiple metrics (see Figure 2A for a schematic). Amplitudes were calculated as the square-root of wavelet-derived power estimates and are used interchangeably throughout the manuscript. The standard measure of window-averaged amplitudes, ***overall amplitudes*** were computed by averaging across the entire segment at its alpha peak frequency. In contrast, ***rhythmic amplitudes*** correspond to the amplitude estimates during detected rhythmic episodes. If no alpha episode was indicated, abundance was set to zero, and amplitude was set to missing. Unless indicated otherwise, both amplitude measures were normalized by subtracting the amplitude estimate of the fitted background spectrum. This step represents a parameterization of rhythmic power (cf. Haller et al., 2018) and is conceptually similar to baseline normalization, without requiring an explicit baseline segment. This highlights a further advantage of rhythm-detection procedures like (e)BOSC. In addition, we calculated an ***overall signal-to-noise ratio***

***(SNR)*** as the ratio of the overall amplitude to the background amplitude: 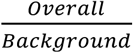. In addition, we defined ***rhythmic SNR*** as the background-normalized rhythmic amplitude as a proxy for the rhythmic representation: 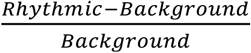.

Unless stated differently, subject-, and condition-specific amplitude and abundance values were averaged within and across trials, and across posterior-occipital channels (P7, P5, P3, P1, Pz, P2, P4, P6, P8, PO7, PO3, POz, PO4, PO8, O1, Oz, O2), in which alpha power was maximal (Figure 4A, Figure 8).

### 2.8 eBOSC validation via alpha rhythm simulations

To assess eBOSC’s detection performance, we simulated 10 Hz sine waves with varying amplitudes (0, 2, 4, 6, 8, 12, 16, 24 [a.u.]) and durations (2, 4, 8, 16, 32, 64, 128, 200 [cycles]) that were symmetrically centred within random 1/f-filtered white noise signals (20 s; 250 Hz sampling rate). Amplitudes were scaled relative to the power of the 8-12 Hz 6^th^ order Butterworth-filtered background signal in each trial to approximate SNRs. To ensure comparability with the empirical analyses, we computed overall SNR analogously to the empirical data, which tended to be lower than the target SNR. We chose the maximum across simulated durations as an upper bound (i.e., conservative estimate) on overall SNR. For each amplitude-duration combination we simulated 500 “trials”. We assessed three different detection pipelines regarding their detection efficacy: the standard BOSC algorithm (i.e., linear background fit incorporating the entire frequency range with no post-editing of the detected matrix); the eBOSC method using wavelet correction by simulating the maximum bias introduced by the wavelet (“MaxBias); and the eBOSC method using the full-width-at-half-maximum amplitude for convolution correction (“FWHM”). The background was estimated separately for each amplitude-duration combination. 500 edge points were removed bilaterally following wavelet estimation, 250 additional samples were removed bilaterally following BOSC detection to account for the duration threshold, effectively retaining 14 s of simulated signal.

Detection efficacy was indexed by signal detection criteria regarding the identification of rhythmic time points between 8 and 12 Hz (i.e., hits = simulated and detected points; false alarms = detected, but not simulated points). These measures are presented as ratios to the full amount of possible points within each category (e.g., hit rate = hits/all simulated time points). For the eBOSC pipelines, abundance was calculated identically to the analyses of empirical data. As no consecutive episodes (cf. Pepisode and abundance) are available in standard BOSC, abundance was defined as the relative amount of time points with detected rhythmicity between 8 to 12 Hz.

A separate simulation aimed at establishing the ability to accurately recover amplitudes. For this purpose, we simulated a whole-trial alpha signal (i.e., duration = 1) and a quarter-trial alpha signal (duration = .25) with a larger range of amplitudes (1:16 [a.u.]) and performed otherwise identical procedures as described above. To assess eBOSC’s ability to disambiguate power and duration (Figure 2B), we additionally performed simulations in the absence of noise across a larger range of simulated amplitudes and durations.

A major change in eBOSC compared to standard BOSC is the exclusion of the rhythmic peak prior to estimating the background. To investigate to what extent the two methods induce a bias between rhythmicity and the estimated background magnitude (for a schematic see Figure 1C and D), we calculated Pearson correlations between the overall amplitude and the estimated background amplitude across all levels of simulated amplitudes and durations (Figure 3C).

**Figure 3:**
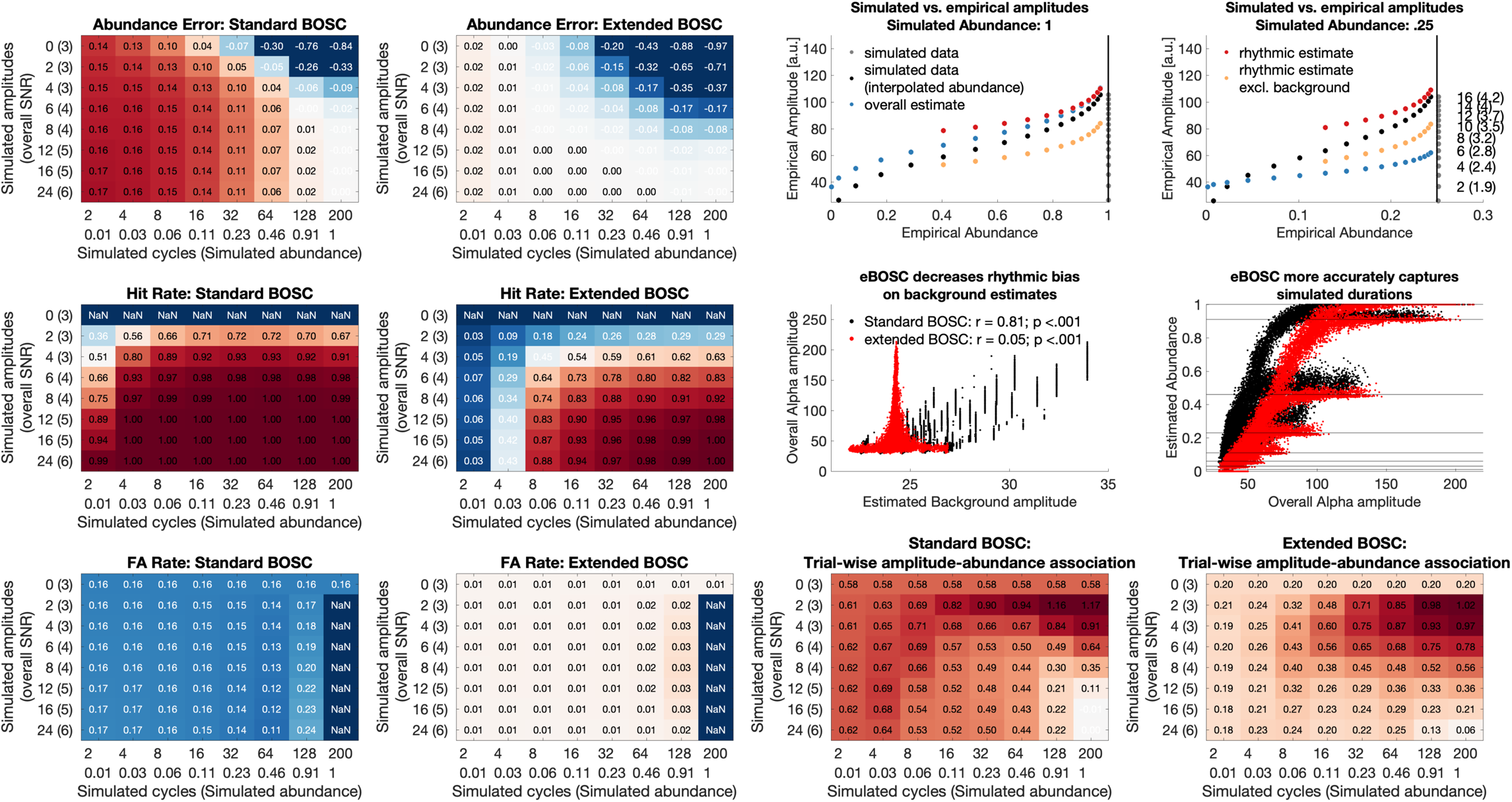
Rhythm detection performance of standard and extended BOSC in simulations. (**A**) Signal detection properties of the two algorithms. For short simulated rhythmicity, abundance is overestimated by standard BOSC, but not eBOSC, whereas eBOSC underestimates the duration of prolonged rhythmicity at low SNRs (A1). Extended BOSC has decreased sensitivity (A2), but higher specificity (A3) compared with extended BOSC. Note that for simulated zero alpha amplitude, all sample points constitute potential false alarms, while by definition no sample point constitutes a potential hit. (**B**) Amplitude and abundance estimate for signals with sustained (left) and short rhythmicity (right). Black dots indicate reference estimates for a pure sine wave without noise, coloured dots indicate the respective estimates for data with the 1/f background. [Note that the reference estimates were interpolated at the empirical abundance of the 1/f data. Grey dots indicate the perfect abundance estimates in the absence of background noise.] When rhythms are sustained (left), impaired rhythm detection at low SNRs causes an overestimation of the rhythmic amplitude. At low rhythmic duration (right), this deficit is outweighed by the severe bias of arrhythmic duration on overall amplitude estimates (e.g., Figure 9). Simulated amplitudes (and corresponding empirical SNRs in brackets) are shown on the right. Vertical lines indicate the simulated rhythmic duration. (**C**) eBOSC successfully reduces the bias of the rhythmic peak on the estimation of the background amplitude. In comparison, standard BOSC induces a strong coupling between the peak magnitude and the background estimate. (**D**) eBOSC indicates abundance more accurately than standard BOSC at high amplitudes (i.e., high SNR; see also A1). The leftward shift indicates a decrease in sensitivity. Horizontal lines indicate different levels of simulated duration. Dots are single-trial estimates across levels of simulated amplitude and duration. (**E**) Standard BOSC and eBOSC induce trial-wise correlations between amplitude and abundance. eBOSC exhibits reduced trial-by-trial coupling at higher SNR compared to standard BOSC. Values are r-to-z-transformed correlation coefficients.

**Figure 4:**
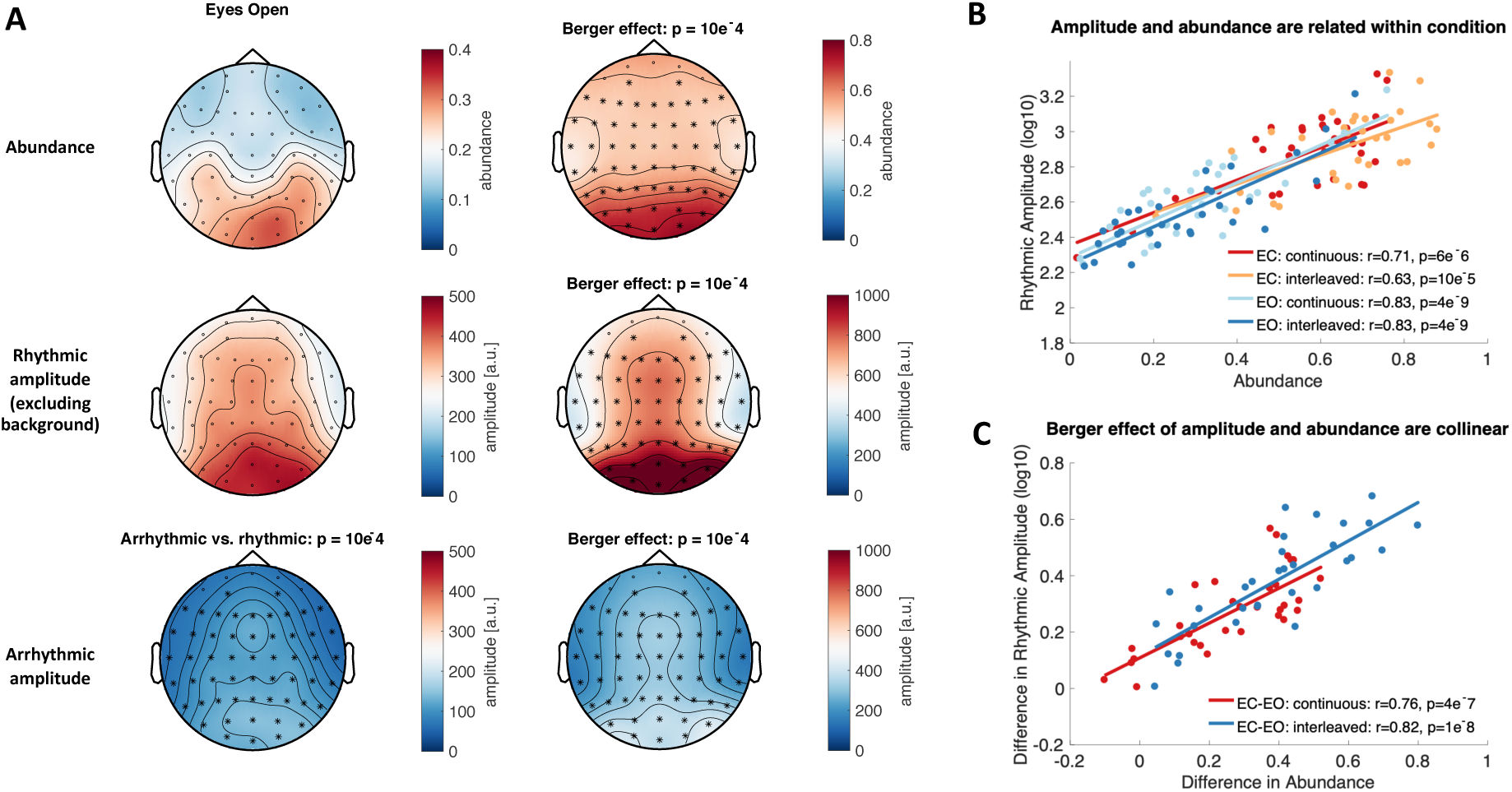
Rhythmic alpha abundance and amplitude during rest. (**A**) eBOSC identifies high occipital alpha abundance and rhythmic amplitude especially during the Eyes Closed resting state. White asterisks indicate significant decreases for arrhythmic from rhythmic amplitudes (cluster is identical between conditions). Black asterisks indicate significant increases upon eye closure. (**B**) Rhythmic amplitude and abundance are inter-individually related during rest (**C**) The modulation of eye closure has similar effects on amplitude and abundance. Estimates were extracted from posterior-occipital channels.

As the empirical data suggested a trial-wise association between amplitude and abundance estimates also at high levels of signal-to-noise ratios (Figure 7), we investigated whether such associations were also present in the simulations. For each pair of simulated amplitude and duration, we calculated Pearson correlations between the overall amplitude and abundance across single trials. Note that due to the stationarity of simulated duration, trial-by-trial fluctuations indicate the bias under fluctuations of the noise background (as amplitudes were scaled to the background in each trial). For each cell, we performed Fisher’s r-to-z transform to account for unequal trial sizes due to missing amplitude/abundance estimates (e.g. when no episodes are detected).

### 2.9 Calculation of phase-based lagged coherence

To investigate the convergence between the power-based duration estimate (abundance) and a phase-based alternative (Fransen et al., 2015), we calculated lagged coherence at 40 linearly scaled frequencies in the range of 1 to 40 Hz for each resting-state condition. Lagged coherence assesses the consistency of phase clustering at a single sensor for a chosen cycle lag (see Fransen et al., 2015 for formulas). Instantaneous power and phase were estimated via 3-cycle wavelets. Data were segmented to be identical to eBOSC’s effective interval (i.e., same removal of signal shoulders as described above). In reference to the duration threshold for power-based rhythmicity, we calculated the averaged lagged coherence using two adjacent epochs à three cycles. We computed an index of alpha rhythmicity by averaging values across epochs and posterior-occipital channels, finally extracting the value at the maximum lagged coherence peak in the 8 to 15 Hz range.

### 2.10 Dynamics of rhythmic probability and rhythmic power during task performance

To investigate the detection properties in the task data, we analysed the temporal dynamics of rhythmic probability and power in the alpha band. We created time-frequency representations as described in section 2.6 and extracted the alpha peak power time series, separately for each person, condition, channel and trial. At the single-trial level, values were allocated to rhythmic vs. arrhythmic time points according to whether a rhythmic episode with mean frequency in the respective range was indicated by eBOSC. These time series were averaged within subject to create individual averages of rhythm dynamics. Subsequently, we z-scored the power time series to accentuate signal dynamics and attenuate between-subject power differences. To highlight global dynamics, these time series were further averaged within- and between-subjects. Figure captions indicate which average was used.

### 2.11 Rhythm-conditional spectra and abundance for multiple canonical frequencies

To assess the general feasibility of rhythm detection outside the alpha range, we analysed the retention interval of the adapted Sternberg task, where the occurrence of theta, alpha and beta rhythms has been reported in previous studies (Brookes et al., 2011; Jensen, Gelfand, Kounios, & Lisman, 2002; Jokisch & Jensen, 2007; Lundqvist et al., 2016; Raghavachari et al., 2001; Tuladhar et al., 2007). For this purpose, we re-segmented the data to cover the final 2 s of the retention interval +-3 s of edge signal that was removed during the eBOSC procedure. We performed eBOSC rhythm detection with otherwise identical parameters to those described in section 2.6. We then calculated spectra across those time points where rhythmic episodes with a mean frequency in the range of interest were indicated, separately for four frequency ranges: 3-8 Hz (theta), 8-15 Hz (alpha), 15-25 Hz (beta) and 25-64 Hz (gamma). We subtracted spectra across the remaining arrhythmic time-points for each range from these ‘rhythm-conditional’ spectra to derive the spectra that are unique to those time points with rhythmic occurrence in the band of interest. For the corresponding topographic representations, we calculated the abundance metric as described in section 2.7 for the apparent peak frequency ranges.

### 2.12 Post-hoc characterization of sustained rhythms vs. transients

Instead of exclusively relying on a fixed *a priori* duration threshold as done in previous applications, eBOSC’s continuous ‘rhythmic episodes’ also allow for a post-hoc separation of rhythms and transients based on the duration of identified rhythmic episodes. This is afforded by our extended post-processing that results in a more specific identification of rhythmic episodes (see Figure 3) and an estimated length for each episode. For this analysis (Figure 10), we set the *a priori* duration threshold to zero and separated the resulting episodes post-hoc based on their duration (shorter vs. longer than 3 cycles) at their mean frequency. That is, any episode crossing the amplitude threshold was retained and episodes were sorted by their ‘transient’ or sustained appearance afterwards. We conducted this analysis in the extended task data to illustrate the temporal dynamics of rhythmic and transient events. To investigate the modulation of rhythm- and transient-specific metrics between the retention phase and the probe phase, we averaged metrics within these two intervals and performed a paired t-test between the two respective intervals for four indices: episode number, duration, frequency and power. Cluster-based permutation tests (Maris & Oostenveld, 2007) as implemented in FieldTrip were performed to control for multiple comparisons. Initially, a clustering algorithm formed clusters based on significant t-tests of individual data points (p <.05; cluster entry threshold) with the spatial constraint of min. three adjacent channels. Then, the significance of the observed cluster-level statistic, based on the summed t-values within the cluster, was assessed by comparison to the distribution of all permutation-based cluster-level statistics. The final cluster p-value that we report in Figures was assessed as the proportion of 1000 Monte Carlo iterations in which the cluster-level statistic was exceeded. Cluster significance was indicated by p-values below .025 (two-sided cluster significance threshold).

### 2.13 Time series representations of detected rhythmic events

To visualize the stereotypic depiction of single-trial rhythmic events, we extracted the time series during individual rhythmic episodes that exceeded a post-hoc duration threshold of three cycles. Individual time series were time-locked to the trough of individual rhythmic episodes and averaged across episodes (Sherman et al., 2016). To avoid unequal sample counts at the edges of episodes, we included additional data padding around the trough prior to averaging. The trough was chosen to be the local minimum during the spectral episode that was closest to the maximum power of the wavelet-transformed signal. To better estimate the local minimum, the time domain signal was low-pass filtered at 25 Hz for alpha and beta, 10 Hz for theta and high-pass-filtered at 20 Hz for gamma using a 6^th^ order Butterworth filter. Filters only served the identification of local minima, whereas unfiltered data were used for plotting. Averaged event dynamics during the first session were visualized for theta at Fz, alpha at O2, beta at FCz and gamma at Fz. To visualize single-trial time-domain signals, we computed moving averages of 150 trials across rhythmic episodes concatenated across all subjects.

We further assessed a potential load-modulation of the rate of rhythmic events during working memory retention by counting the number of individual rhythmic episodes with a mean frequency that fell in a moving window of 3 adjacent center frequencies. This produced a channel-by-frequency representation of spectral event rates, which were the basis for subsequent significance testing using dependent sample regression t-tests and implemented in permutation tests as described in section 2.12.

### 2.14 Modulation of rhythm estimates by working memory load and eye closure

To assess the sensitivity of rhythm-derived indices to experimental manipulations, we compared (1) the effect of eye closure (“Berger effect”) and (2) the effect of working memory load between select rhythm indices. To compare rhythm-specific results with traditional approaches, traditional wavelet estimates were derived using identical parameters as used for eBOSC. We performed confirmatory tests of a parametric increase in posterior alpha power and frontal theta power with memory load based on previous reports in the literature (Jensen et al., 2002; Jensen & Tesche, 2002; Jokisch & Jensen, 2007; Meltzer et al., 2008; Michels, Moazami-Goudarzi, Jeanmonod, & Sarnthein, 2008; Onton, Delorme, & Makeig, 2005; Scheeringa et al., 2009; Tuladhar et al., 2007). In addition, we explored a decrease in frontal theta frequency with load. To reduce the amount of statistical contrasts, we averaged all metrics across sessions before submitting them to statistical tests. Load effects for within-subject trial averages between load conditions were assessed by means of a dependent sample regression t-test, implemented within permutation tests (see section 2.12 for details). Similar cluster-based permutation tests were performed for the effect of eye closure on rhythmic and arrhythmic amplitudes and abundance using a paired samples t-test.

Beyond probing effects on each estimate individually, we probed whether rhythm-specific estimates of duration and magnitude uniquely captured task effects over and above traditional indices. For this purpose, we performed post-hoc linear mixed effects analyses, averaging within the abundance effects clusters. Prior to modelling, values were z-scored across subjects and conditions. In each model, a rhythm-specific index (e.g. abundance) served as the dependent variable, while traditional amplitudes served as a fixed dependent variable. Load or eye closure were modelled as fixed effects with random subject intercepts, assuming compound symmetry. For the load effect, we assessed uniquely explained variance with a post-hoc ANOVA, using marginal sums-of-squares (‘Type III’). Linear mixed effects modelling was performed in R 3.6.1(R, 2019) with the nlme package (Pinheiro et al., 2019).

In addition, we explored effects on theta frequency with cluster-based permutations. To visualize frequency modulations, we performed a post-hoc Fast Fourier Transform (FFT) to specifically characterize rhythmic episodes, while normalizing for their duration. To retain an identical frequency resolution across episodes, we zero-padded episodes of variable duration to a fixed duration of two seconds. We then computed a discrete-time Fourier Transform of individual rhythmic episodes: 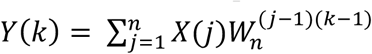, where n is the length of the zero-padded time series X and 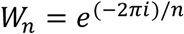, normalized the resulting absolute spectral values by the length of the rhythmic episode *N_rhythmic_* and calculated the single-sided amplitude spectrum. This resulted in rhythm-specific amplitude values with an identical frequency resolution across episodes. In contrast, to derive rhythm-unspecific FFT amplitude estimates, we included the entire two-second retention period in the estimation and used the respective length for normalization, thus resulting in traditional ‘overall’ FFT amplitude estimates that were unspecific to rhythmic occurrence. To assess, whether a theta frequency modulation would be observed with traditional FFT spectra, we detected condition-dependent theta frequency peaks. Peaks were defined as frequencies at which the first derivative of the spectrum changed from positive to negative (Grandy et al., 2013b). In case no peak was identified, the frequency with peak amplitude was selected. Finally, we performed paired-t-tests to estimate potential load effects.

In figures, we display within-subject standard errors (Cousineau, 2005) to highlight condition differences. For these, individual data were centered by subtracting the subject condition average and adding the grand condition average to individual within-condition values.

## 3. Results

### 3.1. Extended BOSC (eBOSC) increases specificity of rhythm detection

We extended the BOSC rhythm detection method to characterize rhythmicity at the single-trial level by creating continuous ‘rhythmic episodes’ (see Figure 1 & Figure S1). A central goal of this approach is the disambiguation of rhythmic power and duration, which can be achieved perfectly in data without background noise (upper row in Figure 2B). However, the addition of 1/f noise reintroduces a partial coupling of the two parameters (lower row in Figure 2B). To better understand the boundary conditions to derive specific amplitude and duration estimates, we compared the detection properties of the standard and the extended (eBOSC) pipeline by simulating varying levels of rhythm magnitude and duration. Considering the sensitivity and specificity of detection, both pipelines performed adequately at high levels of SNR with high hit and low false alarm rates (Figure 3A). However, whereas standard BOSC showed perfect sensitivity above SNRs of ∼4, specificity was lower than for eBOSC as indicated by higher false alarm rates (grand averages: .160 for standard BOSC; .015 for eBOSC). This specificity increase was observed across simulation parameters, suggesting a general abundance overestimation by standard BOSC (see also Figure 3D). In addition, standard BOSC did not show a reduced detection of transient rhythms below the duration threshold of three cycles, whereas hit rates for those transients were clearly reduced with eBOSC (Figure 3A2). This suggests that wavelet convolution extended the effective duration of transient rhythmic episodes, resulting in an exceedance of the temporal threshold. In contrast, by creating explicit rhythmic episodes and reducing convolution effects, eBOSC more strictly adhered to the specified target duration. However, there was also a notable reduction in sensitivity for rhythms just above the duration threshold, suggesting a sensitivity-specificity trade-off (Figure 3A2). In addition to decreasing false alarms, eBOSC also more accurately estimated the duration of rhythmicity (Figure 3A1), although an underestimation of abundance persisted (and was increased) at low SNRs. In sum, while eBOSC improved the specificity of identifying rhythmic content, there were also noticeable decrements in sensitivity (grand averages: .909 for standard BOSC; .614 for eBOSC), especially at low SNRs. Comparable results were obtained with a 3-cycle wavelet (Figure S3). Notably, while sensitivity remains an issue, the high specificity of detection suggests that the estimated rhythmic abundance serves as a lower bound on the actual duration of rhythmicity.

In a second set of simulations, we considered eBOSC’s potential to accurately estimate rhythmic amplitudes. As expected, in signals with stationary rhythms (duration = 1), the time-invariant ‘overall’ amplitude estimate most accurately represented simulated amplitudes (Figure 3B left), as any methods-induced underestimation biased rhythm-specific amplitudes. Specifically, at low SNRs, underestimation of rhythmic content resulted in an overestimation of rhythmic amplitudes, as some low-amplitude time points were incorrectly excluded prior to averaging. At those low SNRs, subtraction of the background estimate (cf. baseline normalization) alleviated this overestimation. The general impairment at low SNRs was however outweighed by the advantage of rhythm-specific amplitude estimates in time series where rhythmic duration was low and thus arrhythmicity was prevalent (Figure 3B right). Here, rhythm-specific estimates accurately tracked simulated amplitudes, whereas a strong underestimation was observed for unspecific power indices. In both scenarios, we observed an underestimation of rhythmic abundance with decreasing amplitudes (cf. Figure 3A1).

An adaptation of the eBOSC method is the exclusion of the rhythmic alpha peak prior to fitting the arrhythmic background. This serves to reduce a potential bias of rhythmic content on the estimation of the arrhythmic content (see Figure 1C for a schematic). Our simulations indeed indicated a bias of the spectral peak amplitude on the background estimate in the standard BOSC algorithm, which was substantially reduced in eBOSC’s estimates (Figure 3C).

To gain a visual representation of duration estimation performance, we plotted abundance against amplitude estimates across all simulated trials, regardless of simulation parameters (Figure 3D). This revealed multiple modes of abundance at high amplitude levels, which in the eBOSC case more closely tracked the simulated duration. This further visualizes the decreased error in abundance estimates, especially at high SNRs (e.g., Figure 3A), while an observed rightward shift towards higher amplitudes indicated the more pronounced underestimation of rhythmicity at low SNRs.

Finally, we investigated the trial-wise association between amplitude and duration estimate based on the observed coupling in empirical data (see Figure 7). Our simulations suggest that both standard BOSC and eBOSC can induce spurious positive correlations between amplitude and abundance estimates, which are most pronounced at low levels of SNR (Figure 3E). Notably, these associations are strongly reduced in eBOSC, especially when rhythmic power is high. This indicates that eBOSC provides a better separation between the two (here independent) parameters, although a spurious association remains.

In sum, our simulations suggest that eBOSC specifically separates rhythmic and arrhythmic time points in simulated data at the expense of decreased sensitivity, especially when SNR is low. However, the increase in specificity is accompanied by an increased accuracy of duration estimates at high SNR, theoretically allowing a more precise investigation of rhythmic duration.

While the simulations provide a gold standard to assess detection performance, we further probed eBOSC’s detection performance in empirical data from resting and task states to investigate the practical feasibility and utility of rhythm detection. As the ground truth in real data is unknown, we evaluated detection performance by contrasting metrics from detected and undetected timepoints regarding their topography and time course.

Individual power spectra showed clear rhythmic alpha peaks for every participant during eyes closed rest and for most subjects during eyes open rest and the task retention period, indicating the general presence of alpha rhythms during the analysed states (Figure S4). In line with a putative source in visual cortex, alpha abundance was highest over parieto-occipital channels during the resting state (Figure 4A) and during the WM retention period (Figure 8), with high collinearity between abundance and rhythmic amplitudes within resting conditions (Figure 4B). As expected, rhythmic time-points exhibited increased alpha power compared with arrhythmic time points (Figure 4A; white cluster). As one of the earliest findings in cognitive electrophysiology (Berger, 1938), alpha amplitudes increase in magnitude upon eye closure. Here, eye closure was reflected by a joint shift towards higher amplitudes and durations for almost all participants (Figure 4C). To assess unique contributions of the Berger effect on rhythm indices while controlling for the high collinearity between indicators, we performed linear mixed modelling within the common effects cluster (see Supplementary Table 1). We focussed on the continuous condition here, due to the similarity of the effects in the interleaved case. Notably, rhythmic abundance was modulated by eye closure while statistically controlling for either rhythmic or arrhythmic amplitudes. In contrast, rhythmic alpha amplitudes were not modulated by eye closure when controlling for alpha abundance. This suggests that rhythmic duration may be a more sensitive marker of task modulations than amplitude. Finally, arrhythmic amplitudes did not exhibit the Berger effect in either the interleaved or the continuous acquisition when statistically controlling for the collinearity with rhythmic amplitude or rhythmic abundance. Taken together, these results suggest a high, joint sensitivity of rhythm-specific indices to eye closure, which exceeded the residual modulation of arrhythmic backgrounds that may have resulted from specificity impairments during the original detection procedure.

The temporal dynamics of indicated rhythmicity are another characteristic of interest to indicate successful rhythm detection. While such an investigation is difficult for induced rhythmicity during rest, evoked rhythmicity offers an optimal test case due to its systematic temporal deployment. For this reason, we analysed task recordings with stereotypic design-locked alpha power dynamics at encoding, retention and probe presentation (Figure 5AB). Rhythmic probability closely tracked power dynamics (Figure 5A) and time points designated as rhythmic exhibited pronounced alpha power compared with those labelled arrhythmic (Figure 5A left vs. Figure 5A right). While rhythm-specific dynamics closely captured standard power trajectories, we observed a dissociation concerning arrhythmic power. Here, we observed transient increases during stimulus onsets that were absent from either abundance or rhythmic power (Figure 5A right). This suggests an increase in high-power transients that were excluded due to the 3 cycle duration threshold. Indeed, a significant increase in transient events was observed without an *a priori* duration threshold (see Figure 10).

**Figure 5:**
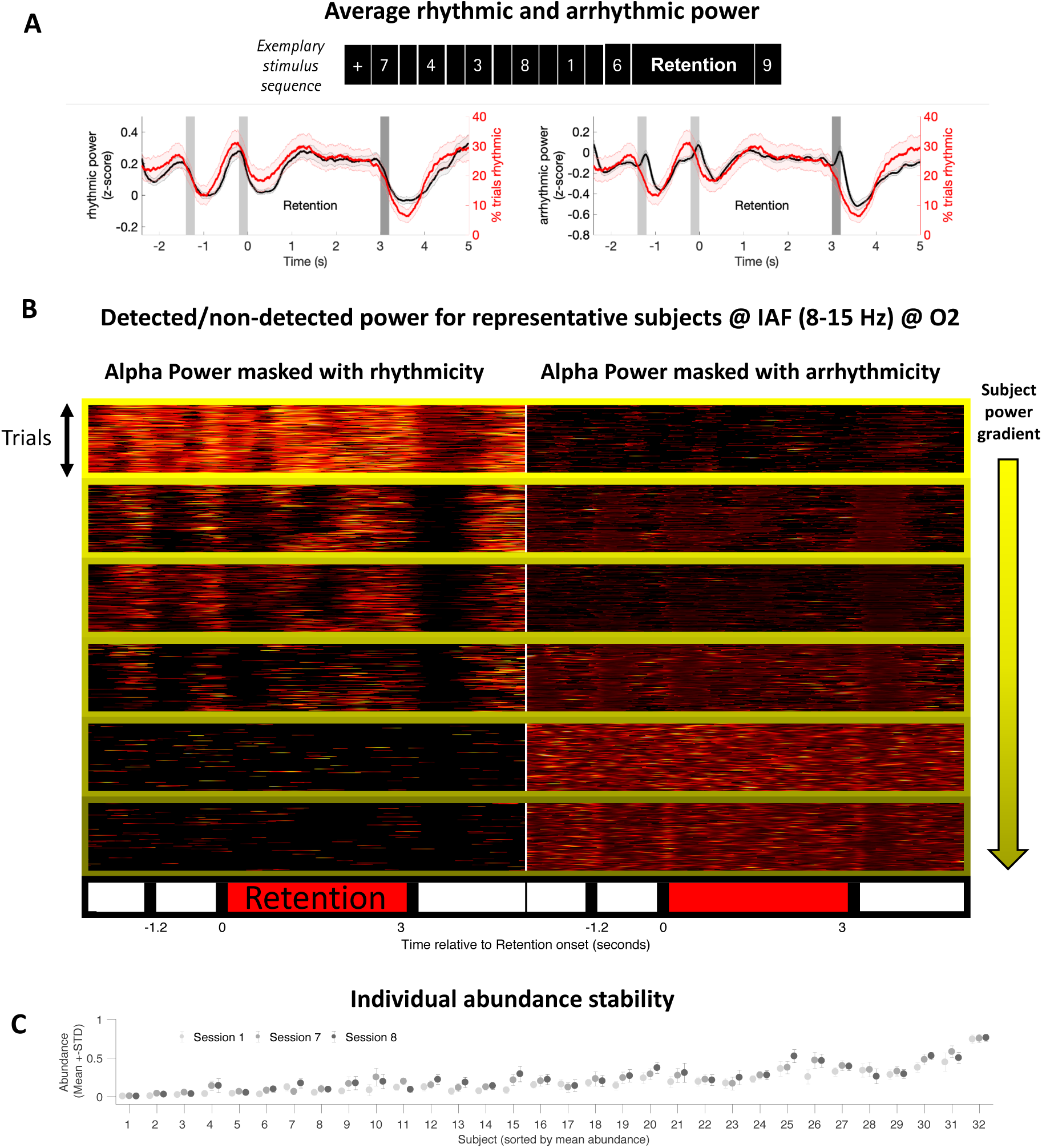
Detected rhythmicity follows the task structure, with stable inter-individual differences in single trial detection. (**A**) Average alpha power (black), split by rhythmic vs. arrhythmic designation, and rhythmic probability (red) at posterior-occipital channels exhibit stereotypic temporal dynamics during encoding (gray bars), retention (0 to 3 s) and retrieval (black bars). Compared to rhythmic power, arrhythmic power exhibits similar temporal dynamics, but is strongly reduced in power (see y-scales). The arrhythmic power dynamics are characterized by additional transient increases following stimulus presentations. Data are from the first session and the high load condition. Shading indicates standard errors across subjects. (**B**) Task-related alpha dynamics are captured by eBOSC at the single-trial level. Each box displays individual trial-wise z-standardized alpha power at the individual peak frequency, separately for rhythmic (left) and non-rhythmic (right) time points. While rhythmic time points (left) exhibit clear single-trial power increases that are locked to the task design, arrhythmic time points (right) do not show evoked task dynamics that separate them from the background, hence suggesting an accurate rejection of rhythmicity. The subplots’ frame colour indicates the subjects’ raw power maximum (i.e., the data scaling). Data are from channel O2 during the first session across load conditions. (**C**) Individual abundance estimates are stable across sessions. Data were averaged across posterior-occipital channels and high (i.e., 6) item load trials.

At the single-trial level, rhythmicity was indicated for periods with visibly elevated alpha power with strong task-locking (Figure 5B left). Conversely, arrhythmicity was indicated for time points with low alpha power and little structured dynamics (Figure 5B right). However, strong inter-individual differences were apparent, with little detected rhythmicity when global alpha power was low (Figure 5B bottom; plots are sorted by descending power as indicated by the frame colour of the depicted subjects and scaled using z-scores to account for global power differences). Crucially, those subjects’ single-trial power dynamics did not present a clear temporal structure, suggesting a prevalence of noise and therefore a correct rejection of rhythmicity. Notably, those individual rhythmicity estimates were stable across multiple sessions (Figure 5C), suggesting that they are indicative of trait-like characteristics rather than idiosyncratic measurement noise (Grandy et al., 2013).

In sum, these results suggest that eBOSC successfully separates rhythmic and arrhythmic episodes in empirical data, both at the group and individual level. However, they also indicate prevalent and stable differences in single-trial rhythmicity in the alpha band that may impair an accurate detection of rhythmic episodes.

### 3.3 Rhythmic SNR constrains empirical duration estimates and rhythm-related metrics

While the empirical results suggest a successful separation of rhythmic and arrhythmic content at the single-trial level, we also observed strong (and stable) inter-individual differences in alpha-abundance. This may imply actual differences in the duration of rhythmic engagement (as indicated in Figure 5B). However, we also observed a severe underestimation of abundance as a function of the overall signal-to-noise ratio (SNR) in simulations (Figure 3), thus leading to the question whether empirical data fell into similar ranges where an underestimation was likely. During the resting state, we indeed observed that many overall SNRs were in the range, where simulations with a stationary alpha rhythm suggested an underestimation of abundance (cf. black and blue lines in Figure 6A. The black line indicates simulation-based estimates for stationary alpha rhythms at different overall SNR levels; see section 2.8). Moreover, the coupling of individual SNR and abundance values took on a deterministic shape in this range, whereas the association was reduced in ranges where simulations suggest sufficient SNR for unbiased abundance estimates (orange line in Figure 6A). As overall SNR is influenced by the duration of arrhythmic signal, rhythmic SNR may serve as an even better predictor of abundance due to its specific relation to rhythmic episodes (Figure 2). In line with this consideration, rhythmic SNR exhibited a strong linear relationship to abundance (Figure 6B). Importantly, the background estimate was not consistently related to abundance (Figure 6C), emphasizing that it is the ‘signal’ and not the ‘noise’ component of SNR that determines detection. Similar observations were made in the task data during the retention phase (Figure S5), suggesting that this association reflects a general link between the magnitude of the spectral peak and duration estimates. The joint analysis of simulated and empirical data thus questions the accuracy of individual duration estimates, especially at low SNRs, due to the dependence of unbiased estimates on sufficient rhythmic power.

**Figure 6:**
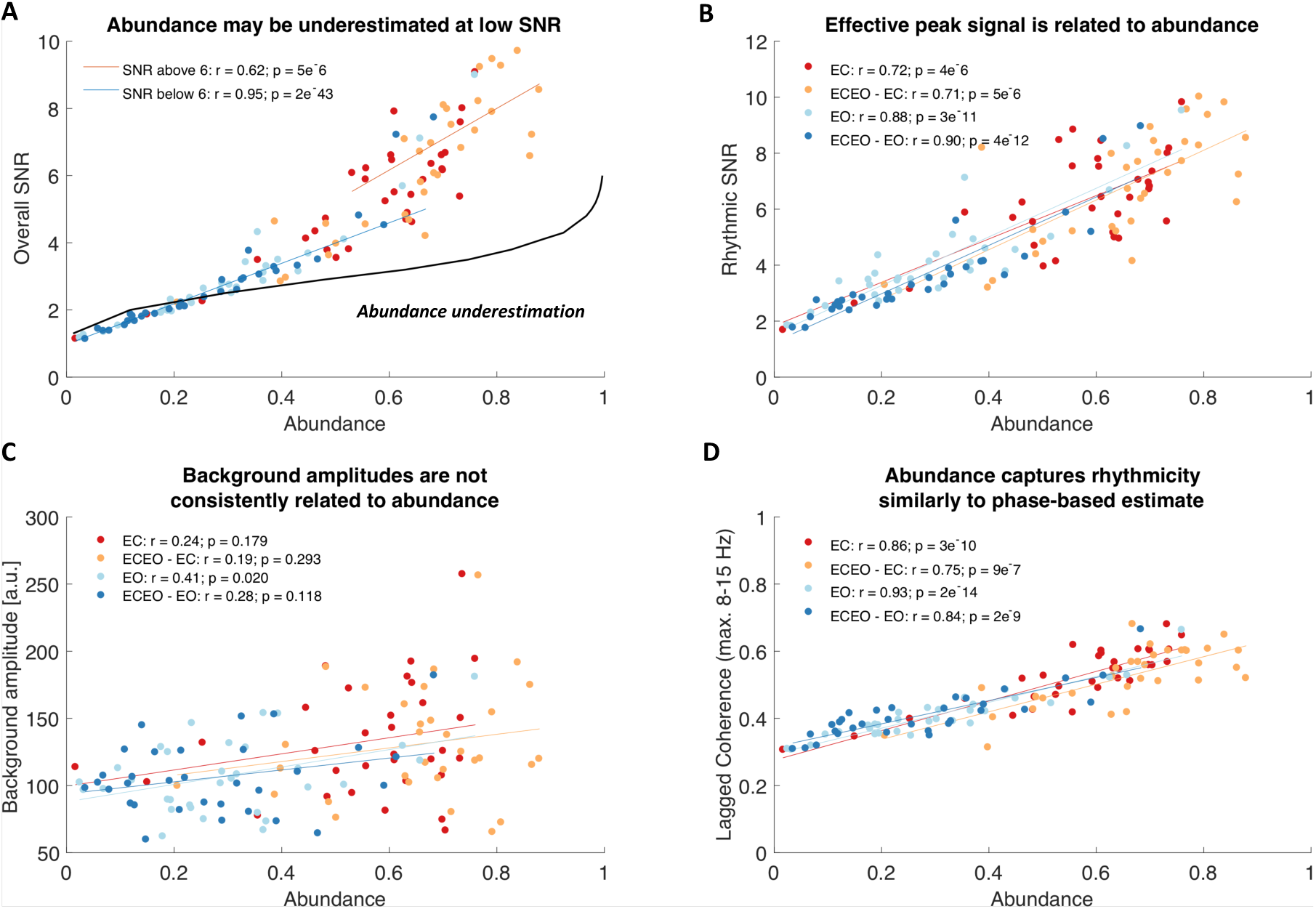
Inter-individual alpha abundance is strongly associated with rhythmic, but not arrhythmic power and may be underestimated at low rhythmic SNR. (**A**) Individual abundance estimates are strongly related to the overall SNR of the spectral alpha peak. This relationship is also observed when only considering individual data within the SNR range for which simulation analyses indicated an unbiased abundance estimation. The black line indicates interpolated estimates from simulation analyses with a sustained rhythm (i.e., duration = 1; see Figure 3B left). Hence, it indicates a lower bound for the abundance underestimation that occurs at low SNRs, with notable overlap with the empirical estimates in the same SNR range. (**B**) The effective rhythmic signal can be conceptualized as the background-normalized rhythmic amplitude above the background estimate (rhythmic SNR). This proxy for signal clarity is inter-individually linked to abundance estimates. (**C**) Background estimates are not consistently related to abundance. This implies that the relationship between amplitude and abundance is mainly driven by the signal, but not background amplitude (i.e., the effective signal ‘clarity’) and that associations do not arise from a misfit of the background. (**D**) Rhythmicity estimates translate between power- and phase-based definition of rhythmicity. This indicates that the BOSC-detected rhythmic spectral peak above the 1/f spectrum contains the rhythmic information that is captured by phase-based duration estimates. All data are from the resting state.

As eBOSC defines single-trial power deviations from a stationary power threshold as a criterion for rhythmicity, it remains unclear whether this association is exclusive to such a ‘power thresholding’-approach or whether it constitutes a more general feature of single-trial rhythmicity. To probe this question, we calculated a phase-based measure of rhythmicity, termed ‘lagged coherence’ (Fransen et al., 2015), which assesses the stability of phase clustering at a single sensor for a chosen cycle lag. Here, 3 cycles were chosen for comparability with eBOSC’s duration threshold. Crucially, this definition of rhythmicity led to highly concordant estimates with eBOSC’s abundance measure^3^ (Figure 6D), suggesting that power-based rhythm detection above the scale-free background overlaps to a large extent with the rhythmic information captured in the phase-based lagged-coherence measure. Moreover, it suggests that duration estimates are more generally coupled to rhythmic amplitudes, especially when overall SNR is low.

While the previous observations were made at the between-subjects level, we further investigated whether such coupling also persists between trials in the absence of between-person differences. In the present data, we indeed observed a positive coupling of trial-wise fluctuations of rhythmic SNR and abundance (mean Fisher’s z: .60; p < 6.5e-19) (Figure 7A), whereas the estimate of the scale-free background was less consistently, though significantly (mean Fisher’s z: .20; p = 2.6e-6), related to the estimated duration of rhythmicity (Figure 7B). This suggests that the level of estimated abundance primarily relates to the magnitude of ongoing power fluctuations around the stationary power threshold. Figure 7C schematically shows how such an amplitude-abundance coupling may be reflected in single trials as a function of rhythmic SNR. These relationships were also observed in our simulations and in other frequency bands, although they were reduced in magnitude at higher levels of simulated empirical SNR (Figure 3E) and for other frequencies (Figure S6), suggesting that partial dissociations of the two parameters are feasible.

In sum, these results strongly caution against the interpretation of duration measures as a ‘pure’ duration metric that is independent from rhythmic power, especially at low levels of SNR. The strong within-subject coupling may however also indicate an intrinsic coupling between the strength and duration of neural synchrony as joint representations of a rhythmic mode. Notably, covariations were not constrained to amplitude and abundance, but were widespread, including covariations between ‘SNR’ and the instability (or variability) of the individual alpha peak frequency (see Supplementary Materials; Figure S7). Combined, these results suggest that the efficacy of an accurate single-trial characterization of neural rhythms relies on sufficient individual rhythmicity and can not only constrain the validity of duration estimates, but broadly affect a range of rhythm characteristics that can be inferred from single trials.

### 3.4 Rhythm detection improves amplitude estimates by removing arrhythmic episodes

From the joint assessment of detection performance in simulated and empirical data, it follows that low SNR constitutes a severe challenge for single-trial rhythm characterization. However, while the magnitude of rhythmicity at the single trial level constrains the detectability of rhythms, abundance represents a lower bound on rhythmic duration due to eBOSC’s high specificity. This allows the interpretation of rhythm-related metrics for those time points where rhythmicity is indicated, leading to tangible benefits over standard analyses. In this section, we highlight multiple proof-of-concept cases of such benefits.

A considerable problem in standard narrowband power analyses is the superposition of rhythmicity on top of a scale-free 1/f background, effectively mixing the two components in traditional power estimates (e.g. Haller et al., 2018). In contrast, eBOSC uncouples the two signals via explicit modelling of the arrhythmic background. Figure 8 presents a comparison between the standard narrowband estimate and eBOSC’s background and rhythmicity metrics for the alpha band during working memory retention. While high narrowband power is observed in frontal and parietal clusters, eBOSC differentiated a frontally-dominated 1/f component and a posterior-occipital rhythm cluster. Identical comparisons within multiple low-frequency ranges suggest the separation of a stationary 1/f topography and spatially varying superpositions of rhythmicity (Figure S8). This highlights a successful separation of the scale-free slope magnitude from rhythmicity across multiple frequencies, even when topographies are partially overlapping as in the case of theta.

Furthermore, the presence of a rhythm is a fundamental assumption for the interpretation of rhythm-related metrics, e.g., phase (Aru et al., 2015). This is often verified by observing a spectral peak at the frequency of interest. However, sparse single-trial rhythmicity may not produce an overt peak in the average spectrum due to the high prevalence of low-power arrhythmic content. Crucially, knowledge about the temporal occurrence of rhythms in the ongoing signal can be used to investigate the spectral content that is specific to those time points, thereby creating ‘rhythm-conditional spectra’. Figure 9A highlights that such rhythm-conditional spectra can recover spectral peaks for multiple canonical frequency bands, even when no clear peak is observed in the grand average spectrum. This showcases that a focus on detected rhythmic time points allows the interpretation of rhythm-related parameters.

**Figure 7:**
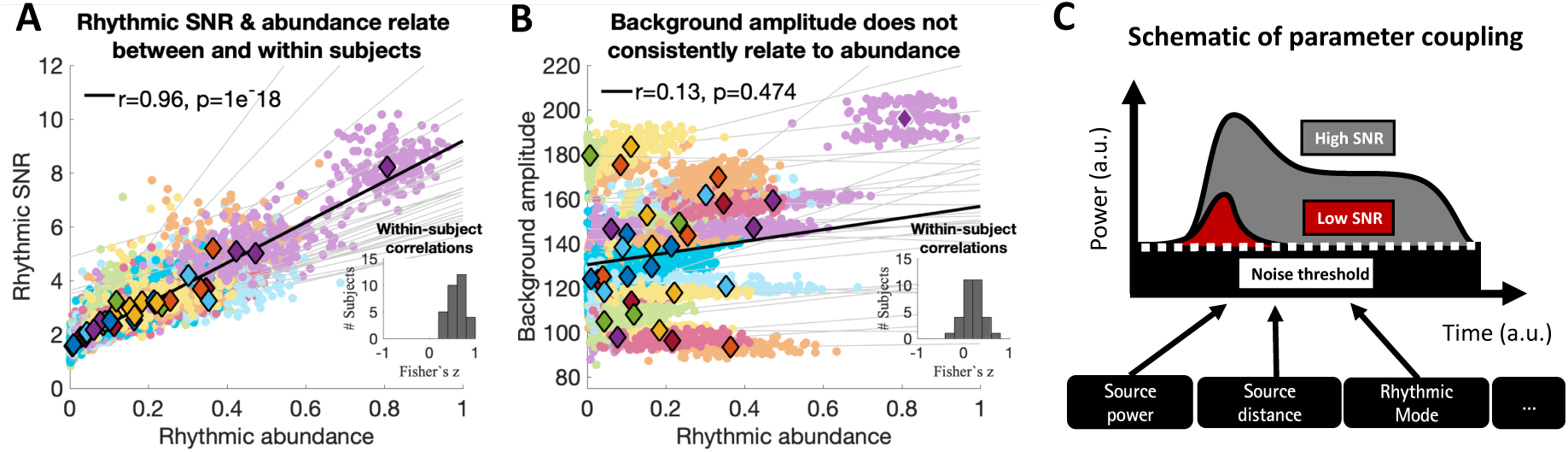
The magnitude and duration of single-trial rhythmicity are intra-individually associated. Amplitude-abundance association within subjects in the Sternberg task (1^st^ session, all trials). Dots represent single trial estimates, color-coded by subject. Subject means are presented via diamonds. (Inlay) Histogram of within-subject Fisher’s z-coefficients of within-subject associations. Relationships are exclusively positive. (B) Background estimates are inter-individually uncorrelated with single-trial abundance fluctuations, excluding the outlier indicated by white edges. (C) Schematic of the potential interdependence of rhythmic SNR and abundance. Low SNR may cause the detection of shorter supra-threshold power periods with constrained amplitude ranges, whereas prolonged periods may exceed the stationary threshold when the rhythmic signal is clearly separated from the background.

**Figure 8:**
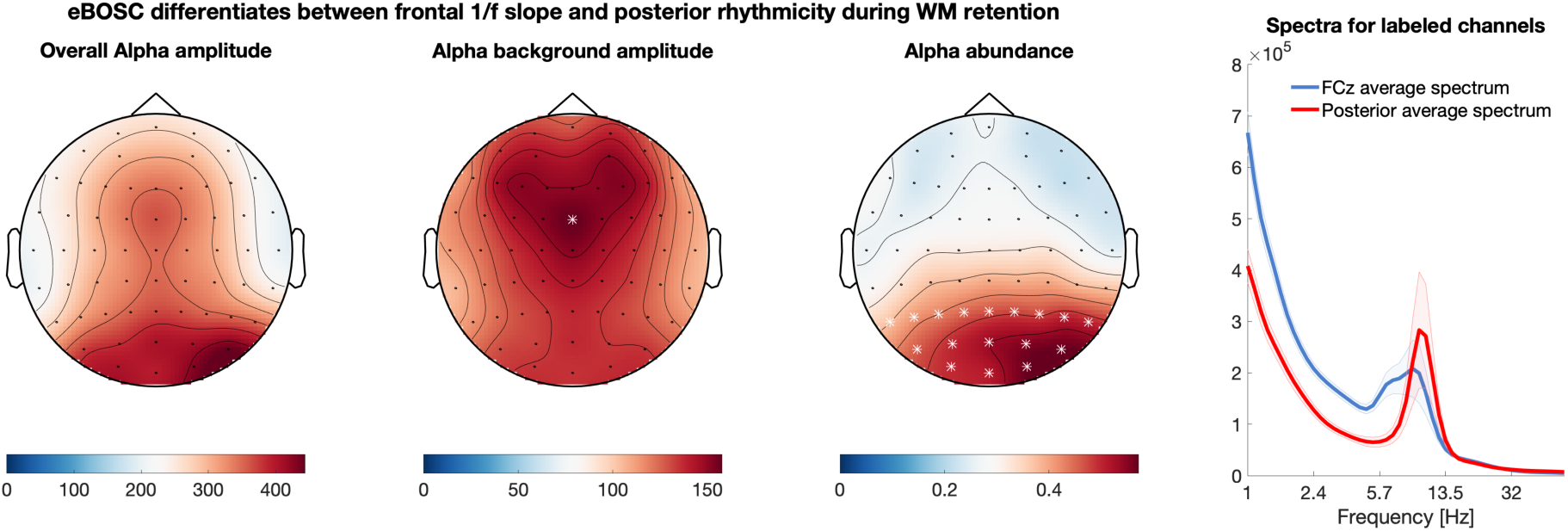
eBOSC differentiates spatially varying topographies of rhythmic and arrhythmic power during working memory retention. Asterisks mark the channels that were selected for the spectra on the right. The graph shading depicts standard errors. The topographies are grand averages from the retention phase of the Sternberg task across all sessions.

**Figure 9:**
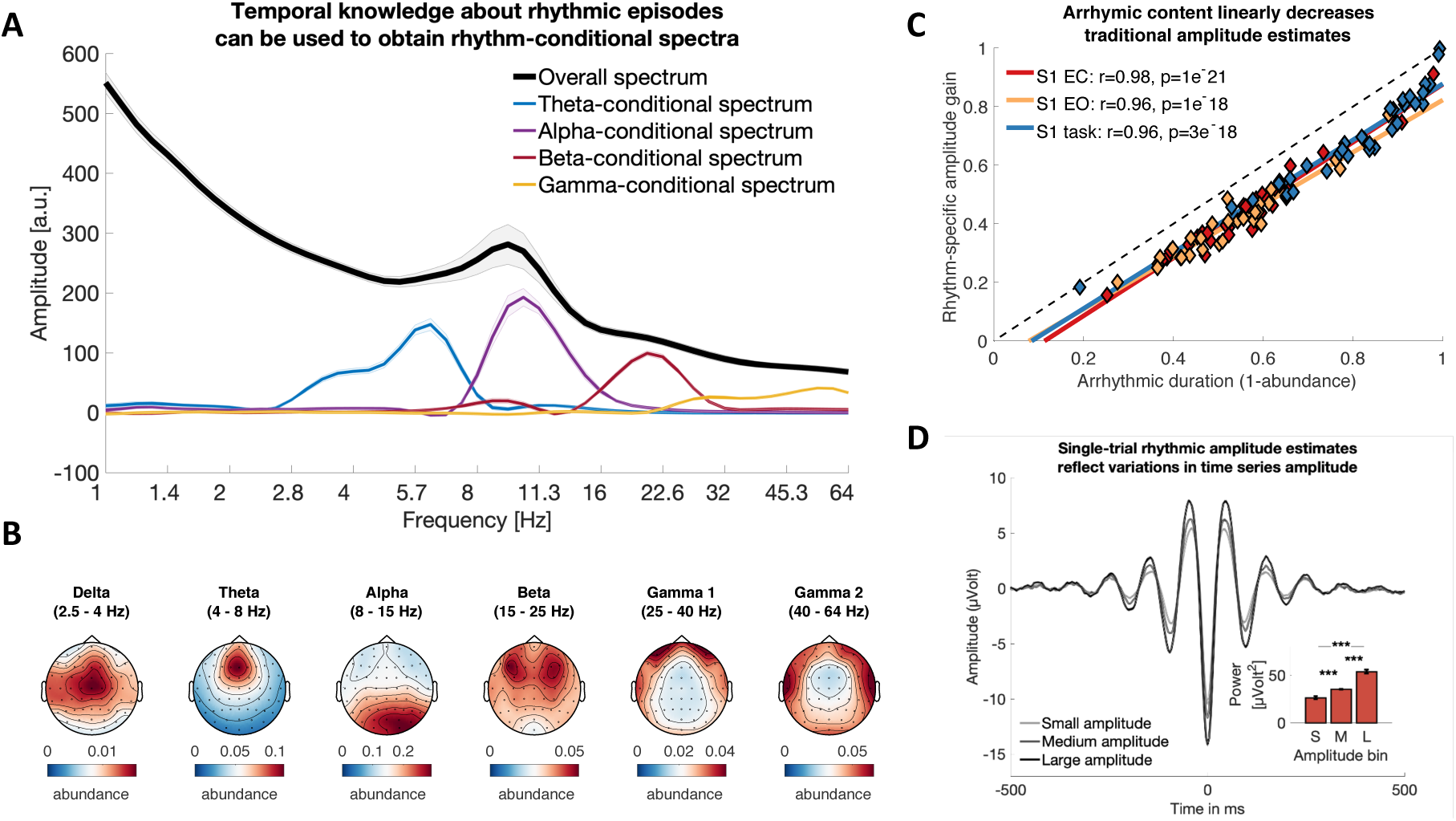
Time-wise indication of rhythmicity improves rhythmic amplitude estimates and produces rhythm-conditional spectra. (A) Comparison of rhythm-conditional spectra with the standard overall spectrum during the memory retention phase. Rhythm-conditional spectra are created by comparing spectra from time-points where a rhythm in the respective frequency range has been indicated with those where no rhythm was present. Notably, this indicates rhythmic peaks at the frequencies of interest that are not observed in the overall spectrum (e.g. theta, beta) due to the prevalence of non-rhythmic events. Simultaneous peaks beyond the target frequencies indicate cross-spectral coupling. Note that these spectra also suggest sub-clusters of frequencies (e.g. an apparent split of the ‘theta-conditional’ spectrum into a putative delta and theta component). Data are averaged across sessions, loads, subjects and channels. (**B**) Abundance topographies of the observed rhythm-conditional spectral peaks. (C) Arrhythmic duration linearly biases traditional power estimates during both rest and task states. The relative gain in alpha amplitudes from global intervals to eBOSC’s rhythmic periods (see schematic in Figure 1A and Figure 2A) increases with the arrhythmic duration in the investigated period. That is, if high arrhythmic duration was indicated, a focus on rhythmic periods strongly increased amplitudes by excluding the pervasive low-amplitude arrhythmic periods. In contrast, amplitude estimates were similar when arrhythmicity was low and hence rhythm-unspecific metrics contained little arrhythmic bias. Dots represent individual condition averages during the resting state. Amplitude gain is calculated as the relative change in rhythmic amplitude from the unspecific ‘overall’ amplitude (i.e., (rhythmic amplitude-overall amplitude)/rhythmic amplitude). (D) Rhythmic amplitudes reflect variations in time series amplitude, here visualized via a triadic split. The inset shows the statistical comparison of squared amplitudes in a 200 ms peri-peak window. Estimates are from Session 1 with data from all channels. *** = p < .001.

Abundance topographies for the different peaks observed in the rhythm-conditional spectra, were in line with the canonical separation of these frequencies in the literature (Figure 9B). Notably, while some rhythmicity was identified in higher frequency ranges, the associated abundance topographies suggests a muscular generator rather than a neural origin for these events.

Related to the recovery of spectral amplitudes from ‘overall amplitudes’, a central prediction of the present work was that the change from overall to rhythmic amplitudes (i.e., rhythm-specific gain; see Figure 2 for a schematic) scales with the presence of arrhythmic signal. Stated differently, if most of the overall signal is rhythmic, the difference between overall and rhythm-specific amplitude estimates should be minimal. Conversely, if the overall signal consists largely of arrhythmic periods, rhythm-specific amplitude estimates should strongly increase from their unspecific counterparts. In line with these expectations, we observed a positive, highly linear, relationship between a subject’s estimated duration of arrhythmicity and the rhythm-specific amplitude gain (Figure 9C). Thus, for subjects with sparse rhythmicity, rhythm-specific amplitudes were strongly increased from overall amplitudes, whereas differences were minute for subjects with prolonged rhythmicity. Note however that in the case of inter-individual collinearity of amplitude and abundance (as observed in the present data) the rhythm-specific gains are unlikely to change the rank-order of subjects as the relative gain will not only be proportional to the abundance, but due to the collinearity also to the original amplitude. While such collinearity was high in the alpha band, decreased amplitude-abundance relationships were observed for other canonical frequency bands (Figure S6), where such ‘amplitude recovery’ may have the most immediate benefits.

To assess whether these single-trial amplitude estimates validly reflected fluctuations in time series magnitude, we performed a triadic split based on single-trial amplitude estimates across all detected episodes (across channels and sessions) in the alpha band. We aligned time-series representations of rhythmicity to the maximal negative peak and compared power in a window of 200 ms around this peak. Notably, rhythm-specific amplitude estimates reflected time series amplitudes during rhythmic periods (Figure 9D) with a larger effect size (medium vs. small: p =4e-7, Cohen’s d = 1.13, large vs. medium: p = 4e-9; Cohen’s d = 1.42) than overall amplitudes (medium vs. small: p =.002, Cohen’s d = .58, large vs. medium: p = 9e-7; Cohen’s d = 1.08). Interestingly, despite collinearity between amplitude and abundance at the within-subject level (Figure 7A), a triadic split based on single-trial abundance estimates did not differentiate rhythmic amplitudes (medium vs. small: p =.34, Cohen’s d = .17, large vs. medium: p = .45; Cohen’s d = -.14). Hence, rhythm-specific amplitude estimates were better predictors of time series amplitudes than traditional averages that included arrhythmic episodes or estimates of rhythmic duration.

In sum, eBOSC provides sensible single-trial amplitude estimates of narrow-band rhythmicity that are boosted in magnitude due to the removal of arrhythmic episodes.

### 3.5 eBOSC separates sustained and transient spectral events

In addition to specificity gains for rhythmic indices, eBOSC’s creation of temporally contiguous rhythmic ’episodes’ affords a characterization of rhythmic and transient episodes with significant spectral power in the absence of an *a priori* duration requirement. Using the traditional 3-cycle threshold as a post-hoc criterion for detected episodes, we separated rhythmic and transient spectral events with clear differences in their time-domain representations (Figure 10A). Notably, while rhythmic SNR related to the number of detected rhythmic events, the same was not observed for the number of transient episodes (Figure 10B2), thus indicating that rhythms and transients may arise from different mechanisms. In line with the observations made for rhythmic vs. arrhythmic power (cf. Figure 5A), we observed differences in the temporal prevalence of transient events and sustained rhythms. Specifically, stimulus onsets increased the number of transient events (Figure 10A1), whereas sustained rhythms were increased during the retention phase. These episodes can be further characterized in terms of their duration in cycles (Figure 10A2), their mean frequency (Figure 10A3) and event-specific power (Figure 10A4). During the retention phase, we observed an increased number of larger and longer rhythms compared with the probe period with no apparent differences in frequency. In contrast, we observed a global increase in the number of transients during probe presentation, with those transients being of higher frequency compared to transients during the retention phase. The magnitude and duration of transients did not differ globally between these two task periods. Taken together, these analyses suggest a principled separation of sustained and transient spectral events on the bases of temporal post-hoc thresholds.

Finally, the temporal specificity of spectral episodes also enables a characterization of rhythm-‘evoked’ events (see Supplementary Materials). Whereas an assessment of evoked effects has thus far only been possible with regard to external event markers, the indication of rhythm on- and offsets allows an investigation of concurrent changes that are time-locked to rhythmic events (Figure S9A). Here, we exemplarily show that the on- and offsets of rhythmic episodes are associated with concurrent power increases and decreases respectively (Figure S9B), adding further evidence for the high temporal specificity of indicated on- and offsets of rhythmic episodes.

In sum, these proof-of-concept applications suggest that explicit rhythm detection may provide tangible benefits over traditional narrowband analyses due to the specific separation of rhythmic and arrhythmic periods, despite the high collinearity of abundance and power that we observed in the alpha band.

### 3.5 Rhythm-specific indices exhibit improved sensitivity to working memory load

So far, we investigated the potential to derive rhythm-specific estimates and highlighted resulting benefits. It remains unclear however, to what extent these estimates are experimentally modulated in cognitive tasks and whether they add complementary information to extant measures. To attend this question, we probed the effect of working memory load on traditional, rhythm-unspecific power averages and eBOSC’s duration and amplitude in the alpha and theta band^4^. Standard power estimates indicated load-related increases in frontal theta and right posterior alpha power that did not reach statistical significance however (Figure 11A; see also Figure S10 for different normalization procedures). In contrast, significant increases were observed for rhythmic abundance (Figure 11B), but not for rhythm-specific power, despite similar statistical topographies (Figure 11C). To investigate whether rhythmic abundance captured additional variance of memory load compared to amplitude, we performed linear mixed effects modeling of data averages within the (topographically-similar) abundance clusters. The results are presented in Supplementary Table 2. As expected, we observed high collinearity between different measures, expressed as significant pairwise relations between traditional and rhythm-specific indices. Controlling for this high collinearity however, memory load predicted increases in theta and alpha abundance over and above overall, and rhythmic-specific, amplitudes. In contrast, rhythm-specific amplitudes did not capture unique variance in load level when controlling for overall amplitude, in line with the absence of an indicated effect by the permutation test. Jointly, these analyses suggest that rhythmic abundance, despite high collinearity with overall and rhythmic amplitudes, is more sensitive to working memory load than (traditional) amplitude estimates.

**Figure 10:**
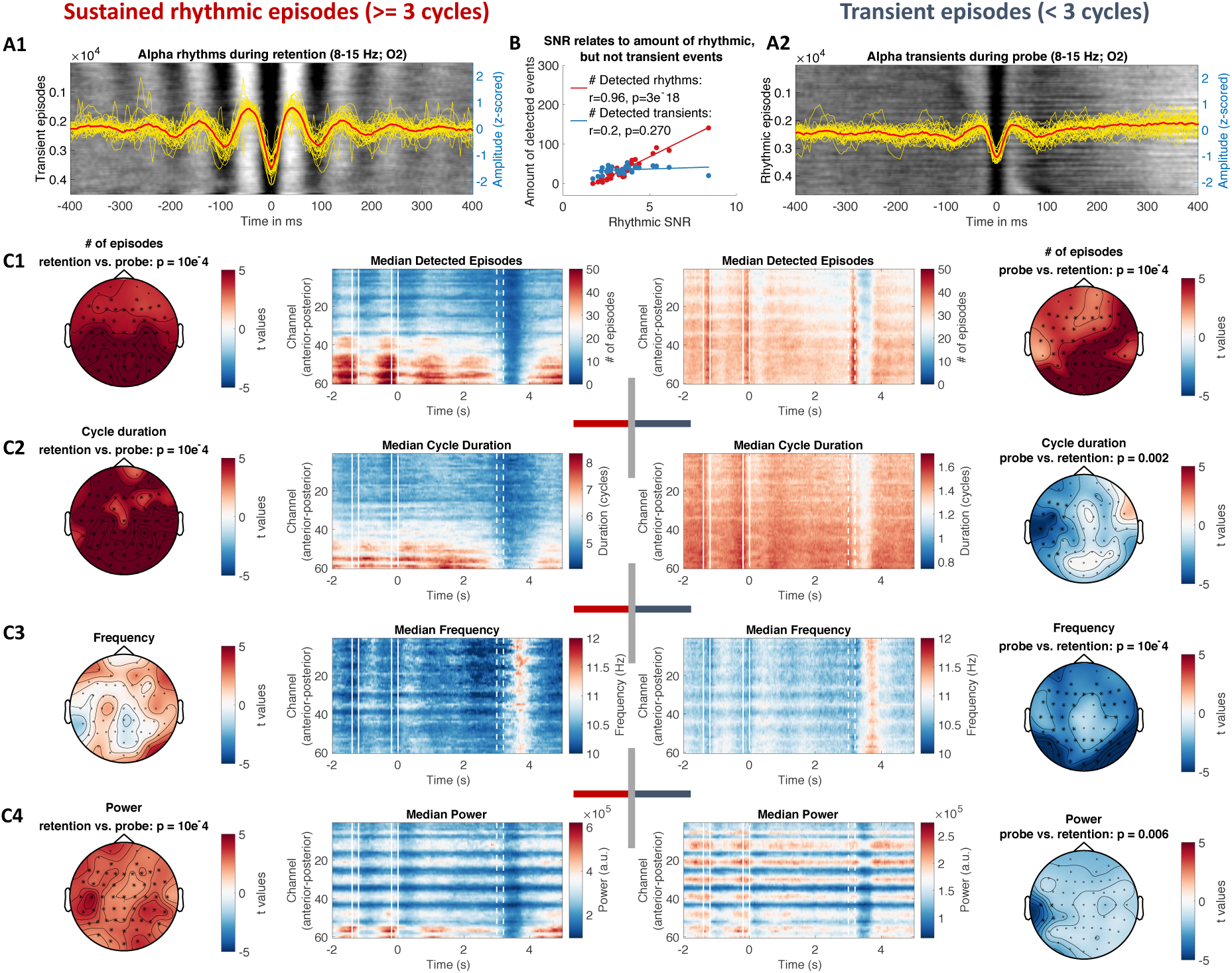
eBOSC provides a varied characterization of duration-specific frequency content, separating sustained rhythmicity from transients. Episodes with a mean frequency between 8 and 15 Hz were post-hoc sorted by falling below or above a 3-cycle duration threshold. For each index, estimates were averaged across all episodes at any time point, followed by averaging across subjects and sessions. All indices are based on episodes that fulfil the power threshold for rhythmicity. (**A**) Time-domain representation of alpha rhythms (A1) and transients (A2) during retention and probe respectively. Backgrounds display moving averages of 150 raw rhythmic episode time series across all subjects. Events are aligned to the closest trough to the TFR maximum of the identified event. Episodes are sorted by episode onset relative to the identified trough. Individual (yellow) and grand data averages (red) are superimposed. (**B**) Rhythmic SNR linearly relates to the number of rhythmic events during retention, but not transient events during probe presentation. (**C**) Rhythm- and transient-specific estimates of episode prevalence (C1), duration (C2), frequency (C3) and power (C4). Central panels show time-channel representations of group indices for rhythmic (left) and transient episodes (right). Lateral topographies indicate the corresponding statistical comparisons of paired t-tests comparing the retention and the probe period. Asterisks signify significant electrode clusters. Unbroken white lines indicate stimulus presentations, broken white lines indicate probe presentation.

**Figure 11:**
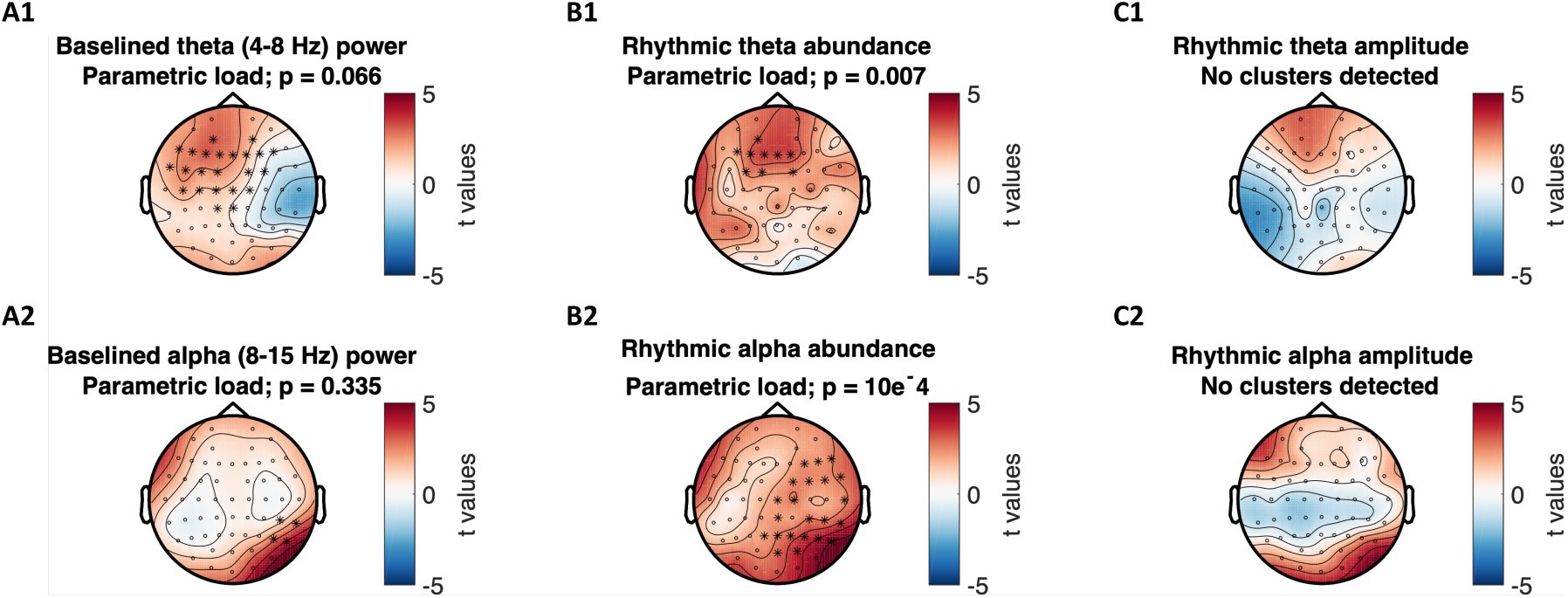
Memory load-modulation of traditional wavelet power, rhythmic abundance and rhythmic amplitude. Traditional wavelet estimates indicated no significant parametric load of either frontal theta or posterior alpha power (A), whereas a load-related increase was indicated for both theta and alpha abundance (B). In contrast to abundance, no significant relationship with load was indicated for rhythm-specific amplitudes (C).

The previous analyses focused on the total rhythmic abundance and power during the retention phase. However, rhythmicity can also be characterized with regard to individual spectral events, such as their rate of occurrence. In line with our observation of high abundance, rhythmic events in the alpha band were characterized by enduring rhythmicity, whereas events in other frequency bands had a more transient signature (Figure 12A). This poses the question whether the rate of these transient events may be a critical parameter, as has been previously suggested for the beta and gamma band (Lundqvist et al., 2016; Shin, Law, Tsutsui, Moore, & Jones, 2017). To attend this question, we created rate spectra based on the occurrence of rhythmic episodes in sliding frequency windows. These spectra were then subjected to a cluster-based permutation test to assess their relation with memory load. We observed increased rates of frontal theta and posterior gamma events as well as decreased rates of central beta events with load, whereas no differences were indicated for the alpha band (Figure 12B). Hence, whereas the sustained appearance of alpha rhythms may render other parameters such as duration and power critical, in other frequency bands, modulation may also affect the number of relatively sparse events.

**Figure 12:**
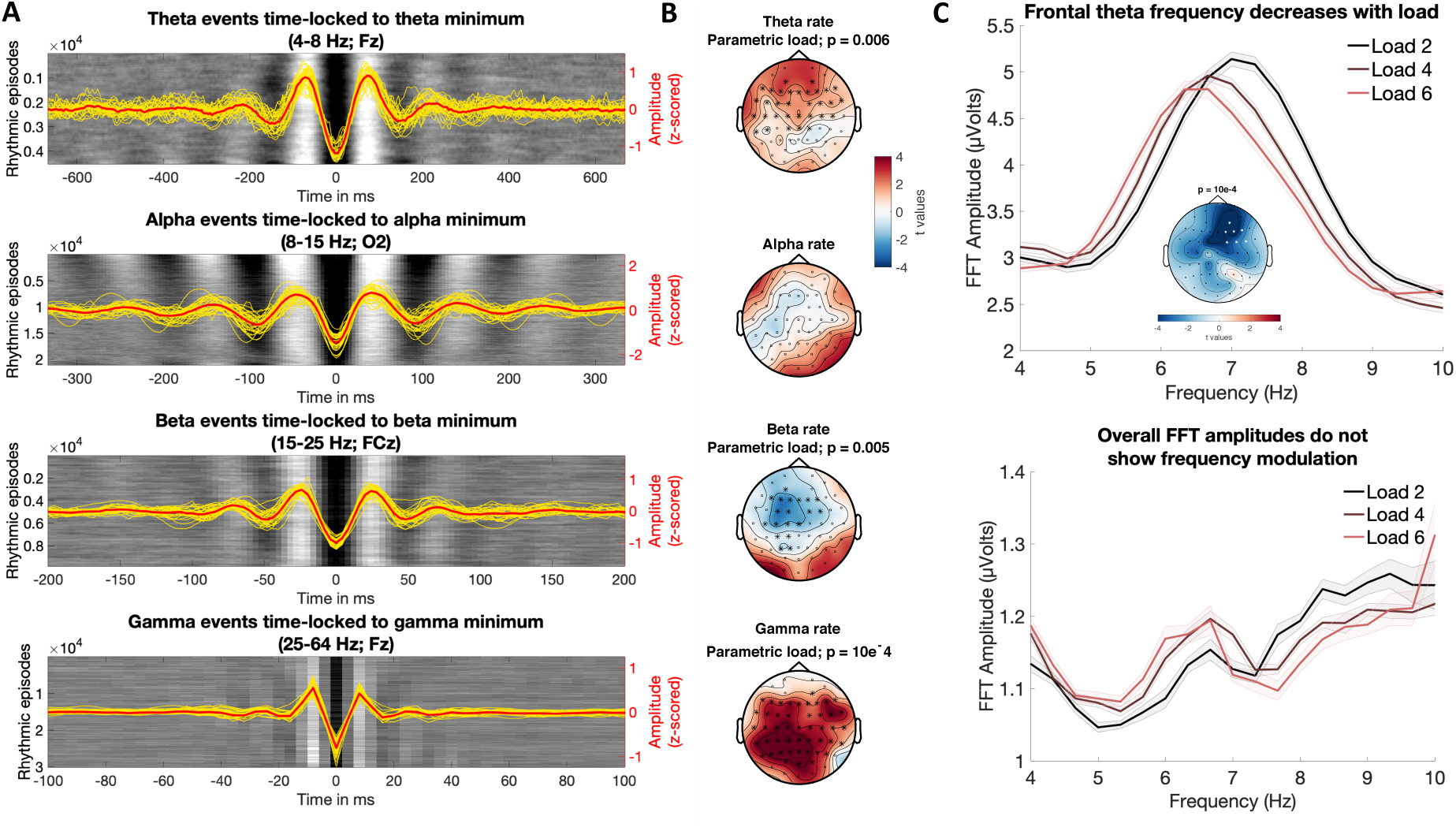
Descriptors of single-trial rhythmic events relate to working memory load. (A, B) Rhythmic event rates are a relevant parameter for describing band-specific task modulations. (A) Different frequency bands vary in their sustained vs. transient time domain appearance. Conventions are the same as in Figure 10B. X-axes are scaled to cover approx. 6 cycles at each frequency. (B) Rhythmic event rates are modulated by working memory load except in the alpha band, where events appear the most sustained. Alpha rate was averaged from 8-12 Hz here to exclude beta rate decreases. (C) Rhythmic frontal theta frequency decreases with working memory load. (Top) Rhythm-specific spectra indicate a parametric shift in theta frequencies with load. Statistics are based on a cluster-based permutation test. The inset shows the cluster for which a significant relation between load and the average frequency of rhythmic theta episodes is indicated. Spectra are averaged across significant cluster channels. Error bars indicate within-subject standard errors. (Lower) The overall spectrum does not show a clear spectral peak in the theta range or a shift in theta frequency. Note that amplitude values are increased in the rhythm-specific version compared to the rhythm-unspecific estimates.

In turn, focusing on these sparse rhythmic events can drastically increase their amplitude estimates and may thus improve dependent metrics (e.g., see Figure 9C). During our exploration of rhythmic parameters, we observed a parametric load-related decrease of frontal theta frequency (Figure 12C) that spatially aligned with the frontal topography of theta rate and abundance increases (see Figure 12B & 11B). Individual rhythmic frequency decreases between low and high loads were not related to individual abundance (r = .33, p = .06) or amplitude (r = .06, p =.73) changes, suggesting that differences in rhythmic SNR cannot solely account for individual frequency shifts. To visualize the shift in theta frequency, we computed FFT spectra with a high spectral resolution (.33 Hz), separately for rhythmic episodes, and – as traditionally done – for the entire retention period. Critically, frequency-modulated theta peaks at frontal channels were only observed for rhythmic, but not for overall spectra (Figure 12C) due to a threefold increase in the magnitude of single-trial events across the entire segment. Moreover, in line with the results of eBOSC’s wavelet-based frequency estimates, significant negative load-related slopes were indicated for rhythm specific FFT frequency estimates (mean= -.16, SE = .05, p = .005) but not rhythm-unspecific global estimates (mean = -.05, SE = .06, p = .36). Hence, a focus on rhythmic episodes was necessary to reveal memory-load related frequency decreases of frontal theta rhythms, which would have been missed with traditional analyses.

In sum, these results highlight the potential of single-trial-based rhythm estimates to boost signal of interest to advance analyses regarding the role of rhythmicity in cognition.

## 4. Discussion

In the present manuscript, we explored the feasibility of characterizing neural rhythms at the level of single trials. To achieve this goal, we extended a previously published rhythm detection method, BOSC (Whitten et al., 2011). Based on simulations we demonstrate that our extended BOSC (eBOSC) algorithm performs well and increases detection specificity. Crucially, the reliance on robust regression in conjunction with removal of the rhythmic power band effectively decoupled estimation of the noise background from the rhythmic signal component (as reflected in the divergent associations with rhythmicity estimates). In real data, we can successfully separate rhythmic and arrhythmic, sometimes transient components, and further characterize e.g., their amplitude, duration and frequency. In total, single-trial characterization of neural rhythms appears promising for improving a mechanistic understanding of rhythmic processing modes during rest and task.

However, the simulations also reveal challenges for accurate rhythm characterization in that the abundance estimates clearly depend on rhythmic power. The comparison to a phase-based rhythm detection further suggests that this a general limitation independent of the chosen detection algorithm. Below, we will discuss the potential and challenges of single-trial rhythm detection in more detail.

### 4.1 The utility and potential of rhythm detection

Single-trial analyses are rapidly gaining importance (Jones, 2016; Stokes & Spaak, 2016), in part due to a debate regarding the sustained vs. transient nature of neural rhythms that cannot be resolved at the level of data averages (Jones, 2016; van Ede et al., 2018). In short, due to the non-negative nature of power estimates, time-varying transient power increases may be represented as sustained power upon averaging, indicating an ambiguity between the duration and power of rhythmic events (cf., Figure 2B). Importantly, sustained and transient events may differ in their neurobiological origin (Sherman et al., 2016), indicating high theoretical relevance for their differentiation. Moreover, many analysis procedures, such as phase-based functional connectivity, assume that estimates are directly linked to the presence of rhythmicity, therefore leading to interpretational difficulties when it is unclear whether this condition is met (Aru et al., 2015; Muthukumaraswamy & Singh, 2011). Clear identification of rhythmic time periods in single trials is necessary to resolve these issues. In the current study, we extended a state-of-the-art rhythm detection algorithm, and systematically investigated its ability to characterize the power and duration of neural alpha rhythms at the single-trial level in scalp EEG recordings.

While the standard BOSC method provides a sensible detection of rhythmic activity in empirical data (Caplan et al., 2015; Whitten et al., 2011), its’ ability to detect rhythmicity and disambiguate rhythmic power and duration has not yet been investigated systematically. Furthermore, we introduced multiple changes that aimed to create rhythmic episodes with a time-point-wise indication of rhythmicity. For these reasons, we assessed the performance of both algorithms in simulations. We observed that both algorithms were able to approximate the duration of rhythmicity across a large range of simulated amplitudes and durations. However, standard BOSC systematically overestimated rhythmic duration (Figure 3A). Furthermore, we observed a bias of rhythmicity on the estimated background (Figure 3C) as also noted by Haller et al. (2018). In contrast, eBOSC accounts for these problems by introducing multiple changes: First, by excluding the rhythmic peak prior to fitting the arrhythmic background, eBOSC decreased the bias of narrow-band rhythmicity on the background fit (Figure 3C), thereby effectively uncoupling the estimated background amplitude from the indicated rhythmicity. Second, the post-processing of detected segments provided a more specific characterization of neural rhythms compared to standard BOSC. In particular, accounting for the temporal extension of the wavelet increased the temporal specificity of rhythm detection as indicated by a better adherence to the *a priori* duration threshold along with more precise duration estimates (Figures 3). In contrast to the high specificity, the algorithm did trade off sensitivity, leading to sensitivity losses that were most pronounced at low signal-to-noise ratios (SNR). In sum, the simulations highlight that eBOSC provides a sensible differentiation of rhythmic and arrhythmic time points as well as accurate duration estimates, but also highlight challenges for empirically disentangling rhythmic power and duration that arise from sensitivity problems when the magnitude of rhythms is low. We discuss this further in section 4.2. In empirical data, eBOSC likewise led to a sensible separation of rhythmic from arrhythmic topographies (Figure 4A, Figure 8, Figure S8) and time courses, both at the average (Figure 5A) and the single-trial level (Figure 5B). This suggests a sensible separation of rhythmic and arrhythmic time points also in empirical scenarios.

The specific separation of rhythmic and arrhythmic time points has multiple immediate benefits that we validated using empirical data from resting and task states. First, eBOSC separates the scale-free background from superimposed rhythmicity in a principled manner. The theoretical importance of such separation has previously been highlighted (Haller et al., 2018), as narrow-band estimates traditionally confound the two signals. Here, we show that such a separation empirically produces different topographies for the arrhythmic background and the superimposed rhythmicity (Figure 8 and Figure S8). In line with these findings, Caplan et al. (2015) described a rhythmic occipital alpha topography, whereas overall power included an additional anterior component across multiple lower frequencies. While that study did not plot topographies for the background estimates, our study suggests that this frontal component is captured by the background magnitude. This provides convergent evidence for a principled separation of rhythmic and arrhythmic spectral content which may be treated as a signal of interest in itself (Buzsáki & Mizuseki, 2014; He et al., 2010).

The separation of these signal sources at single time points can further be used to summarize the rhythmic single-trial content via rhythm-conditional spectra (Figure 9). Crucially, such a focus on rhythmic periods resolves biases from arrhythmic periods in the segments of interest. In line with our hypotheses, simulations (Figure 2B) and empirical data (Figure 9C) indicate that arrhythmic episodes in the analysed segment bias overall power estimates relative to the extent of their duration. Conversely, a focus on rhythmic periods induces the most pronounced amplitude gains when rhythmic periods are sparse. This is in line with previous observations by Cole & Voytek (2018), showing dissociations between power and frequency estimates when considering ‘rhythmic’ vs. unspecific periods and extend those observations by showing a strong linear dependence between the rhythm-specific change in estimates and the duration of arrhythmic bias (Figure 9C).

Moreover, by allowing a post-hoc duration threshold, eBOSC can disentangle transient and sustained events in a principled manner (Figure 10). This may provide new insights into the contribution of different biophysical signal generators (Sherman et al., 2016) to observed neural dynamics and aid the characterization of these processes. Such characterization includes multiple parameters, such as the frequency of rhythmic episodes, their duration, their amplitude and other indices that we did not consider here (e.g., instantaneous phase, time domain shape). Here, we observed an increased number of alpha transients following stimulus onsets, and more sustained rhythms when no stimulus was presented (Figure 5A, Figure 10). In line with these observations, Peterson & Voytek (2017) recently proposed alpha ‘bursts’ to increase visual gain during stimulus onsets and contrasted this role with decreased cortical processing during sustained alpha rhythms. Our data supports such a distinction between sustained and transient events, although it should be noted that the present transients resemble single time-domain deflections that are resolved at alpha frequency (Figure 10A2) and may therefore not directly relate to the ‘rhythmic bursts’ proposed by Peterson & Voytek (2017). Note that the reported duration of ‘burst’ events in the literature is still diverse, often exceeding the 3-cycle threshold used here (Peterson & Voytek, 2017). In contrast to eBOSC however, previous work has not accounted for the impact of wavelet duration. It is thus conceivable that power transients that were previously characterized as 3 cycles or longer are actually shorter after correcting for the impact of wavelet convolution, as is done in the current eBOSC implementation (Figure S1). This temporal specificity also allows an indication of rhythm-evoked changes, here exemplified with respect to rhythm-evoked power changes (Figure S9). We observed a precise and systematic time-locking of power changes to the on- and offset of detected rhythmic episodes. This further validates the detection assumptions of the eBOSC method (i.e. significant power increases from the background), and highlights the temporal specificity of eBOSC’s rhythmic episodes.

In total, eBOSC’s single-trial characterization of neural rhythms provides multiple immediate benefits over traditional average-based analyses temporally precise indication of rhythmic and arrhythmic periods. It thus appears promising for improving a mechanistic understanding of rhythmic processing modes during rest and task.

### 4.2 Single-trial detection of rhythms: rhythmic SNR as a central challenge

The aforementioned examples highlight the utility of differentiating rhythmic and arrhythmic periods in the ongoing signal. However, the simulations also indicated problems to accurately do so when rhythmic power is low. That is, the recognition of rhythms was more difficult at low levels of SNR, leading to problems with their further characterization. In particular, our simulations suggest that estimates of the duration (Figure 6A) and frequency stationarity (Figure S7) increasingly deviate from the simulated parameters as the SNR decreases. Changes in instantaneous alpha frequency as a function of cognitive demands have been theorized and reported in the literature (Haegens, Cousijn, Wallis, Harrison, & Nobre, 2014; Herrmann, Murray, Ionta, Hutt, & Lefebvre, 2016; Mierau, Klimesch, & Lefebvre, 2017; Samaha & Postle, 2015; Wutz, Melcher, & Samaha, 2018), with varying degrees of control for power differences between conditions and individuals. Our empirical analyses suggest an increased trial-by-trial variability of individual alpha frequency estimates as SNR decreases (Figure S7). Meanwhile, simulations suggest that such increased variance - both estimated within indicated rhythmic periods and across whole trials – may result from lower SNR. While our results do not negate the possibility of real frequency variations of the alpha rhythm with changes in task load, they emphasize the importance of controlling for the presence of rhythms, mirroring considerations for the interpretation of phase estimates (Muthukumaraswamy & Singh, 2011) and amplitudes. This exemplifies how stable inter-individual differences in rhythmicity (whether due to a real absence of rhythms or prevalent measurement noise; e.g., distance between source and sensor; head shape; skull thickness) can affect a variety of ‘meta‘-indices (like phase, frequency, duration) whose estimation accuracy relies on apparent rhythmicity.

The challenges for characterizing rhythms with low rhythmic power also apply to the estimated rhythmic duration, where the issue is particularly challenging in the face of legitimate interest regarding the relationship between the power and duration of rhythmic events. In particular, sensitivity problems at low rhythmic magnitudes challenge the ability to empirically disambiguate rhythmic duration and power, as it makes the former dependent on the latter in the presence of noise (e.g., Figure 2B). Crucially, a tight link between these parameters was also observed in the empirical data. During both rest and task states, we observed gradual and stable inter-individual differences in the estimated extent of rhythmicity that were most strongly related to the overall SNR in ranges with a pronounced sensitivity loss in simulations (see Figure 4A black line). Given the observed detection problems in our simulations, this ambiguates whether low empirical duration estimates indicate temporally constrained rhythms or estimation problems. Conceptually, this relates to the difference between lower SNR subjects having (A) low power, transient alpha engagement or (B) low power, sustained alpha engagement that was too faint to be detected (i.e., sensitivity problems). While the second was the case in the simulations, the absence of a ground truth does not allow us to resolve this ambiguity in empirical data.

Empirically, multiple results suggest that the low duration estimates at low SNRs did not exclusively arise from idiosyncrasies of our algorithm. Notably, inter-individual differences in eBOSC’s abundance measure were strongly correlated with standard BOSC’s Pepisode measure (Whitten et al., 2011) as well as the phase-based lagged coherence index (Fransen et al., 2015), thus showing high convergence with different state-of-the-art techniques (Figure 6D). Furthermore, detection performance was visually satisfying in single trials given observable task-locked rhythm dynamics for rhythmic, but not arrhythmic periods (Figure 5B). Moreover, the observed relationship between amplitude gain and abundance suggests a successful exclusion of (low-power) arrhythmic episodes at the individual level (Figure 9C). These observations indicate that low SNR conditions present a fundamental challenge to single-trial characterization across different methods. The convergence between power- and phase-based definitions of rhythmicity also indicates that rhythmicity can exhaustively be described by the spectral peak above the background, in line with our observations regarding rhythm-conditional spectra (Figure 9A).

The observation of strong between-person coupling as a function of SNR suggests that such sensitivity limitations may account for the inter-individual amplitude-abundance associations. However, we also observed a positive association between subjects with high alpha SNR. Likewise, we observed positive associations between abundance and rhythmic SNR at the within-subject level (Figure 5). While trial-wise coupling was also present in our simulations, the magnitude of these relationships was lower at high SNR (Figure 3E). Conversely, in empirical data, the within-subject association did not vary in magnitude as a function of the individual SNR. Hence, separate sources may contribute to a coupling of rhythmic amplitude and abundance: a methods-induced association in low SNR ranges and an intrinsic coupling between rhythmic strength and duration as a joint representation of rhythmic synchrony. Notably, empirical within-subject coupling between rhythmic amplitude and duration was previously described for LFP beta bursts in the subthalamic nucleus (Tinkhauser et al., 2017), with both parameters being sensitive to a drug manipulation. This association was interpreted as a “progressive synchronization of inputs over time” (Tinkhauser et al., 2017; p. 2978). Due to the absence of a dissociation of these parameters, it remains unclear whether the two measures make independent contributions or whether they can be conceptualized as a single underlying latent ‘rhythmicity’ index. To resolve this ambiguity, clear dissociations of amplitude and duration estimates in data with high rhythmic SNR are necessary. Notably, potential dissociations between the individual power and duration of beta events has been suggested by Shin et al. (2017), who described differential relationships between event number, power and duration to mean power and behaviour.

The high collinearity between overall amplitude and abundance may be surprising given evidence of their potential dissociation in the case of beta bursts (where overall abundance is low, but burst amplitudes are high) (Lundqvist et al., 2016; Sherman et al., 2016; Shin et al., 2017). In line with this notion, Fransen et al. (2015) reported an increased sensitivity for central beta rhythmicity using the lagged coherence duration index compared with overall power. It may thus be that the alpha range is an outlier in this regard due to the presence of relatively sustained rhythmicity (Figure 12A). A frequency-wise comparison of the between- and within-subject collinearity between amplitude and abundance collinearity indicated a particularly high overlap for the alpha range (Figure S6) with relatively lower coupling for delta, theta and beta. In addition, we observed load modulations on rhythm event rate in many bands but alpha (Figure 12B). Whether these band-specific differences primarily relate to their lower rhythmicity in the current data or reflect systematic differences between frequencies remains an open question and requires data with more prominent rhythmicity in these bands.

The strong collinearity of amplitude and duration estimates also questions the successful disambiguation of the two indices in empirical data and more generally the interpretation of duration as an independent index. In cases where such metrics only serve as a sensitive and/or specific replacement for power (Caplan et al., 2015; Fransen et al., 2015) this may not be problematic, but care has to be taken in interpreting available duration indices as power-independent characteristics of rhythmic episodes. An independent duration index becomes increasingly important however to assess whether rhythms are stationary or transient. For this purpose, both amplitude thresholding and phase-progression criteria have been proposed (Cole & Voytek, 2018; Peterson & Voytek, 2017; Sherman et al., 2016; van Ede et al., 2018; Vidaurre, Myers, Stokes, Nobre, & Woolrich, 2018). Here, we show that both methods arrive at similar conclusions regarding individual rhythmic duration and that the mentioned challenges are therefore applicable to both approaches. As an alternative to threshold-based methods, Van Ede et al. (2018) propose methods based on e.g., Hidden Markov Models (Vidaurre et al., 2018; 2016) for the estimation of rhythmic duration. These approaches are interesting as the definition of states to be inferred in single trials is based on individual (or group) averages, while the multivariate nature of the signals across channels is also considered. It is a viable question for future investigations whether such approaches can adequately characterize the duration of rhythmic states in scenarios where the present methods fail.

### 4.3 Experimental manipulation of rhythm-specific indices

To establish the practical utility of rhythm detection, we probed the experimental modulation of rhythm-specific indices during working memory retention. We focused on this phase as it has received large interest for distinguishing between transient and sustained retention codes (Lundqvist et al., 2016; Lundqvist, Herman, Warden, Brincat, & Miller, 2018), with both theoretical models (Jensen & Lisman, 1998; Lisman & Jensen, 2013; Lundqvist, Herman, & Lansner, 2011) and empirical evidence (Jensen et al., 2002; Jensen & Tesche, 2002; Jokisch & Jensen, 2007; Meltzer et al., 2008; Michels et al., 2008; Onton et al., 2005; Scheeringa et al., 2009; Tuladhar et al., 2007) suggesting that low-frequency rhythmicity increases with load. In line with this evidence, we observed load-related increases in the total duration of frontal theta and right parietal alpha rhythms during visual working memory retention, despite traditional power estimates not reaching statistical significance. Reinforcing these results, mixed modelling indicated a high sensitivity of rhythmic abundance to both eye closure and working memory load while controlling for its collinearity with traditional estimates. This may be due to multiple advantages: eBOSC’s estimates are spectrally normalized and individually specific e.g. to individual peak frequencies, while not assuming stationarity across time. Furthermore, rhythm-specific measures are theoretically agnostic to the magnitude of desynchronization, as they only characterize rhythmicity when it is present. Interestingly, abundance was also more sensitive to the load effect than rhythm-specific amplitudes, suggesting that duration may be a critical parameter to describe cognitive effects despite high collinearity with amplitude.

In addition to our confirmatory analyses in the theta and alpha band, we also explored the load modulation of individual spectral events. Here, we observed that the rate of spectral events during the retention phase was modulated in the theta, beta and gamma, but not the alpha band. This is interesting given that alpha events had a more continuously ‘rhythmic’ appearance overall, whereas the relative rate of spectral events may be relevant for frequency bands with sparse events, as has been suggested for the beta band (Shin et al., 2017). While we confirm the feasibility of such analyses across multiple frequency bands here, we note that further work on the complementary value of such event rates is required to establish their functional significance.

During our analyses we also observed frequency decreases of rhythmic episodes in the theta band at frontal channels. Decreases in rhythmic theta frequency have previously been hypothesized in the framework of theta-gamma multiplexing serving working memory storage (Bahramisharif, Jensen, Jacobs, & Lisman, 2018; Jensen & Lisman, 1998). In particular, a version of this computational model anticipates that the frequency of theta rhythms determines the amount of gamma cycles that can be multiplexed within a single theta cycle. As the number of targets to be held in memory increases, the theory predicts a slowing of theta with increasing load. Such a load-related decrease in gamma-modulating theta frequencies has been observed in human hippocampus (Axmacher et al., 2010). However, this has been difficult to show outside of invasive recordings. Here we observed that overall power did not exhibit a clear spectral peak in the theta range, but that such peak became apparent only when estimates were constrained to rhythmic periods. Furthermore, a parametric decrease in the frequency of single-trial rhythmic episodes was indicated. This suggests that the observed frontal theta signature may support the multiplexing of individual items during the retention period and may even have a hippocampal origin. However, as we observed this effect by exploration, further work should confirm these hypotheses.

Taken together, our results highlight that a variety of rhythm-specific characteristics are sensitive to experimental modulations, such as working memory load. Despite the observed high collinearity between estimates, modulations suggest sensitivity differences between different rhythm estimates. Their automatic single-trial estimation using tools such as eBOSC may thus further our understanding of the role of rhythmicity in cognition, without necessitating the (often unchecked) assumptions of data averages.

### 4.4 Comparison to other single-trial detection algorithms & limitations

The BOSC-family of methods is conceptually similar to other methods that are currently used to identify and describe spectral events in single trials. These methods share the underlying principle of identifying rhythmic events based on momentary power increases relative to an average baseline. Such detection is most common regarding transient beta bursts, for which a beta-specific power threshold is often defined. For example, Sherman et al. (2016) identified transient beta events based on the highest power within the beta range, i.e., without an explicit threshold. Shin et al. (2017) introduced a beta-specific power threshold based on average pre-stimulus power. Similarly, Feingold et al. (2015) defined beta events as exceeding 1.5/3 times the median beta power of that channel, while Tinkhauser et al. (2017) applied a 75^th^ percentile threshold to beta amplitudes. These approaches therefore use a spectrally local power criterion, but no duration threshold. Most closely related to the BOSC-family is the MODAL method by Watrous et al. (2018), which similarly uses a robust fit of the 1/f spectrum to detect rhythmic events in continuous data and then further derives frequency and phase estimates for those rhythmic periods. This is conceptually similar to eBOSC’s definition as ‘statistically significant’ deviations in power from the 1/f background spectrum, except for the absence of a dedicated power or duration threshold. However, all of the above methods share the fundamental assumption of a momentary power deviation from a frequency-specific ‘background’, with varying implementations of a 1/f model assumption. Such assumption can be useful to avoid a bias of rhythmic content on the power threshold (as a spectrally local power threshold depends on the average magnitude of band-limited rhythmicity, i.e., arrhythmic + rhythmic power). Removing the rhythmic peak prior to background modelling helps to avoid such bias (Figure 3C). The eBOSC method thereby provides a principled approach for the detection of single-trial events across frequencies (as shown in Figure 9).

A systematic and general removal of spectral peaks remains a challenge for adequate background estimates. In the current application, we exclusively removed alpha-band power prior to performing the background fit. While the alpha rhythm produced the largest spectral peak in our data (see Figure S4), this should not be understood as a fixed parameter of the eBOSC approach, as other rhythmic peaks may bias the estimation of the background spectrum depending on the recording’s specifics (e.g., type, location etc.). We perceive the need to remove rhythmic peaks prior to background fitting as a general one^5^, as residual spectral peaks bias detection efficacy across the entire spectrum via misfits of the background intercept and/or slope. In particular, rhythmic peaks at higher frequencies disproportionally increase the background estimate at lower frequencies due to the fitting in logarithmic space. Thus, a principled removal of ***any*** spectral peaks in the average spectrum is necessary. Recently, Haller et al. (2018) proposed a principled approach for the removal of rhythmic spectral peaks, which may afford rhythm-unbiased background estimates without requiring priors regarding the location of spectral peaks. It may thus represent a useful pre-processing step for further applications. Regarding the present data, we anticipate no qualitative changes compared to our alpha exclusion approach as (a) we did not consistently observe an association between background and rhythmicity estimates (Figure 6), and the signal was dominated by an alpha frequency peak, which consistently exceeded eBOSC’s power threshold.

Our results further question the adequacy of a stationary power threshold (as traditionally employed and used here) for assessing the amplitude-duration relationship between individual rhythmic episodes. In our empirical analyses, the rhythmic SNR, reflecting the deviation of amplitudes during rhythmic periods from the stationary background, was consistently most strongly associated with the estimated duration (Figures 6 & 7). While keeping the background (and thus the power threshold) stable conforms with the common assumption of rhythmicity being captured within a spectral peak deviating from a stationary background (Figure 9), it may also exacerbate an amplitude-abundance coupling on a trial-by-trial basis (see Figure 7C for a schematic of the assumed association) as ongoing power fluctuations can only be explained by changes in the rhythmic and not the arrhythmic power term. Further research on dynamic thresholds may shed further light on this issue.

Another point worth highlighting is that eBOSC operates on wavelet-derived power estimates. The specific need for wavelet estimates results from model-based assumptions about the time-frequency extension of the wavelet that are used for refining detected rhythmic time points (see Figure 2 and section 2.6). Naturally, the choice of wavelet parameters, specifically their center frequency and duration, influences the time-frequency representations upon which eBOSC operates. Here, we used 6 cycles as the duration parameter, in line with previous work with standard BOSC (Caplan et al., 2015; Whitten et al., 2011). In a supplementary analysis, we compared detection performance using a 3 cycle wavelet and found increased accuracy only for short rhythmicity, whereas the sensitivity to longer rhythmicity was decreased (Figure S3). This is consistent with the assumption that wavelet duration regulates the trade-off between temporal and spectral specificity, with longer wavelets allowing for a finer separation of nearby frequencies at the cost of temporal specificity. Another free parameter concerns the choice of center frequencies. In the post-processing procedures, we perform a sort of spectral filtering based on the pass-band of the wavelet (Figure S1), which is determined by its duration. Resolving rhythms at nearby frequencies thus requires the use of wavelets with sufficient frequency resolution, not only with regard to the sampled frequencies, but also a sufficient duration of the wavelet. This highlights the dependence of eBOSC outputs on the specifics of the wavelet-based transformation from the time into the frequency domain.

An alternative, parallel approach to characterize ongoing rhythmicity is based on characterizing the waveform shape in the time domain, thereby circumventing power analyses entirely (Cole & Voytek, 2018). While such an approach is intriguing, further work is needed to show which analysis sequence is more fruitful: (a) identifying events in the frequency domain and then describing the associated waveform shape in the time domain (e.g., eBOSC) or (b) identifying events and characterizing them based on time domain features (e.g., cycle-by-cycle analysis). As both procedures operate on the basis of single trials, similar challenges (i.e., especially rhythmic SNR) are likely to apply to both approaches.

## 5. Conclusion

We extended a state-of-the-art rhythm detection method and characterized alpha rhythms in simulated, resting and task data at the single trial level. By using simulations, we show that rhythm detection can be employed to derive specific estimates of rhythmicity, with fine-grained control over its definition, and to reduce the bias of rhythm duration on amplitude estimates that commonly exists in standard analysis procedures. However, we also observe striking inter-individual differences in the indicated duration of rhythmicity, which for subjects with low alpha power may be due to insufficient single-trial rhythmicity. We further show that low rhythmicity can lead to biased estimates, in particular underestimated duration and increased variability of rhythmic frequency. Given these constraints, we have provided examples of eBOSC’s efficacy to characterize rhythms that may prove useful for investigating the origin and functional role of neural rhythms in health and disease, and in turn, the current study works to establish the foundation for ideographic analyses of neural rhythms.

## Data availability

The scripts implementing the eBOSC pipelines are available at github.com/jkosciessa/eBOSC alongside the simulation scripts that were used to assess eBOSC’s detection properties. Data will be made available upon reasonable request.

## Acknowledgements

We thank our research assistants and participants for their contributions to the present work. We thank our anonymous reviewers and Scott R. Cole for their helpful comments on earlier versions of this manuscript.

## Footnotes

1 This procedure is similar to calculating the background spectrum from conditions with attenuated alpha power (e.g., the eyes open resting state; Caplan, Bottomley, Kang & Dixon (2015)). However, here we ensure that alpha power is sufficiently removed, whereas if conditions with reduced alpha peak magnitudes are selected, alpha power may still remain sufficiently elevated to influence slope or intercept estimates. Furthermore, the reliance on conditions with decreased rhythmicity appears less suitable given inter-individual differences in alpha engagement in e.g., the eyes open condition. This may induce an implicit contrast to eyes open rhythmicity. Note that when the frequency range is chosen so that the alpha peak represents the middle of the chosen interval, the alpha-induced bias would be captured by a linear increment in the intercept of the background fit, which may also be alleviated by choosing a higher percentile for the power threshold. Notably, removing the alpha peak as done here attenuates such bias, even in cases where the alpha peak biases the slope of the background fit, as would happen if the alpha peak is not centered within the range of sampled frequencies.

2 When multiple alpha-band peaks are present or the peak has a broader appearance, the spectral peak may not be removed entirely, which could result in misfits of the background spectrum. For this purpose, we employed robust regression to down-weight potential residuals around the alpha peak. Our current implementation only accounts for a peak in the alpha range, but could be extended to other frequency ranges using the same logic (see discussion on limitations in section 4.6).

3 The eBOSC duration measure was further strongly correlated with the traditional Pepisode measure (estimated at the trial-wise IAF) that results from the standard BOSC algorithm (EC: r = .96, p = 2e-18; EC2: r = .94, p = 2e-15; EO: r = .97, p = 3e-20; EO2: r = .97, p = 2e-20), suggesting that both measures are similarly sensitive in our empirical data and reflect to a large extent overlapping information.

4 Regarding traditional metrics, we assessed three normalization procedures: raw signals, single-trial log10-transformation and baseline correction with average power 700 to 500 ms prior to retention onset. In contrast with temporal baselining, eBOSC performs spectral normalization by explicitly modelling the 1/f slope.

5 A potential bias is less likely in the case of sporadic rhythmicity that does not produce a peak in the average spectrum. In this case, the power of the single-trial events would exceed the background estimate that is decreased due to the prevalence of arrhythmic periods.

