## Supplementary Information for "Single-trial characterization of neural rhythms: potential and challenges"

**This PDF file includes:**

Supplementary text
Figures S1 to S10
Tables S1 to S2

### SI Methods

**Rhythmic frequency variability during rest.** As an exemplary characteristic of rhythmicity, we assessed the stability of IAF estimates by considering the variability across trials of the task as a function of indicated rhythmicity. Trial-wise rhythmic IAF variability (Figure S7A) was calculated as the standard deviation of the mean frequency of alpha episodes (8-15 Hz). That is, for each trial, we averaged the estimated mean frequency of rhythmic episodes within that trial and computed the standard deviation across trials. Whole-trial IAF variability (Figure S7B) was similarly calculated as the standard deviation of the IAF, with single-trial IAF defined as the frequency with the largest peak magnitude between 8-15 Hz, averaged across the whole trial, i.e., encompassing segments both designated as rhythmic and arrhythmic. Finally, we compared the empirical variability with that observed in simulations (see section 2.8).

**Depiction of rhythm-evoked effects.** The temporal specificity of rhythmic episodes further allows the assessment of 'rhythm-evoked' effects in the temporal or spectral domain. Here, we showcase the rhythm-evoked changes in the same frequency band to indicate the temporal specificity of the indicated rhythmic periods (Figure S9). For this purpose, we calculated time-frequency representations (TFRs) using 6-cycle wavelets and extracted power in the theta (3-8 Hz), alpha (8-15 Hz), beta (15-25 Hz) and gamma-band (25-64 Hz) in 2.4 s periods centred on the on- and offset of indicated rhythmic periods in the respective band. Separate TFRs were calculated for the detected episodes in each channel, followed by averaging across episodes and channels. Finally, we z-transformed the individual averages to highlight the consistency across subjects.

### SI Results

**IAF variability varies as a function of abundance.** Given the strong dependence of accurate duration estimates on sufficient rhythmic power, we investigated how the differences in rhythmicity affect the single-trial estimation of another characteristic, namely the individual alpha frequency (IAF) that generally shows high temporal stability (i.e., trait-qualities) within person at the average level (Grandy, Werkle-Bergner, Chicherio, Schmiedek, et al., 2013b) We observed a strong negative association between the estimated rhythmicity and fluctuations in the rhythmic IAF between trials (Figure S7A). That is, for subjects with pervasive alpha rhythms, IAF estimates were reliably stable across trials, whereas frequency estimates varied when rhythmicity was low. Notably a qualitatively and quantitatively similar association was observed in simulations with a stationary alpha frequency (black lines in Figure S7), suggesting that such variation may be artefactual. As lower abundance implies a smaller number of samples from which the IAF is estimated, this effect could amount to a sampling confound. However, we observed a similar link between overall SNR and IAF variability when the latter was estimated across all timepoints in a trial (Figure S7B). Simulations with stationary 10 Hz rhythms gave rise to similar results, suggesting that estimated frequency fluctuations can arise (at least in part) from the absence of clear rhythmicity. Hence, even when the IAF is intra-individually stable, its moment-to-moment estimation may induce variability when the rhythms are not clearly present.

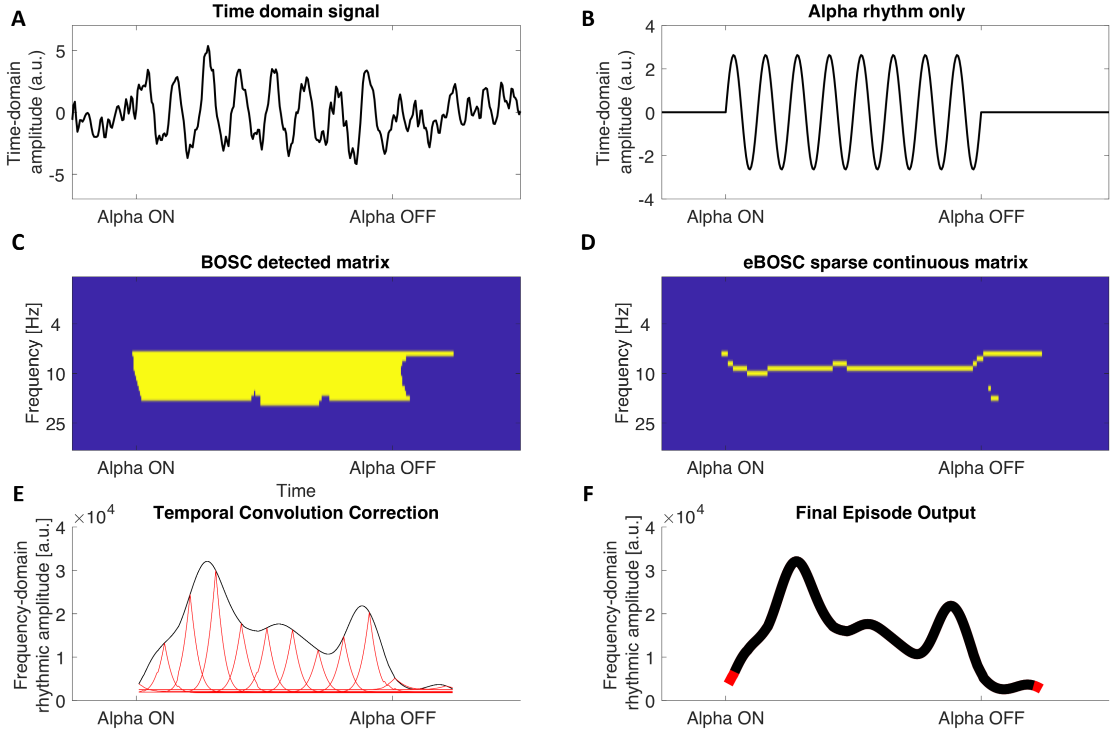


**Fig. S1.** Example of eBOSC's post-processing routines to derive sparse continuous rhythmic 'episodes'. (A) Simulated signal containing 1/f noise and superimposed 10 Hz rhythmicity. (B) 10 Hz rhythmic signal only. (C) Traditional output of BOSC detection: a binary matrix indicates time-frequency points that adhere to power and duration thresholds (in yellow). These matrices are used to calculate *Pepisode*. (D) First step of eBOSC's post-processing: the detected matrix is 'sparsified' in the spectral dimension to create continuous rhythmic episodes. (E) Second step of eBOSC's post-processing: each episode is temporally corrected for the temporal wavelet convolution by estimating the bias of each time point on adjacent time points (here exemplified for select time points via red traces). Only time points that exceed the bias estimated from surrounding time points are retained. (F) Example of final episode trace. The black line indicates the time points that were retained, whereas the red segments were removed during step E. The final episode output is then characterized according to e.g., mean frequency, duration and amplitude, whereas the time points of rhythmicity can for example be used to define rhythm-conditional spectra. These episodes are used to calculate *abundance*.

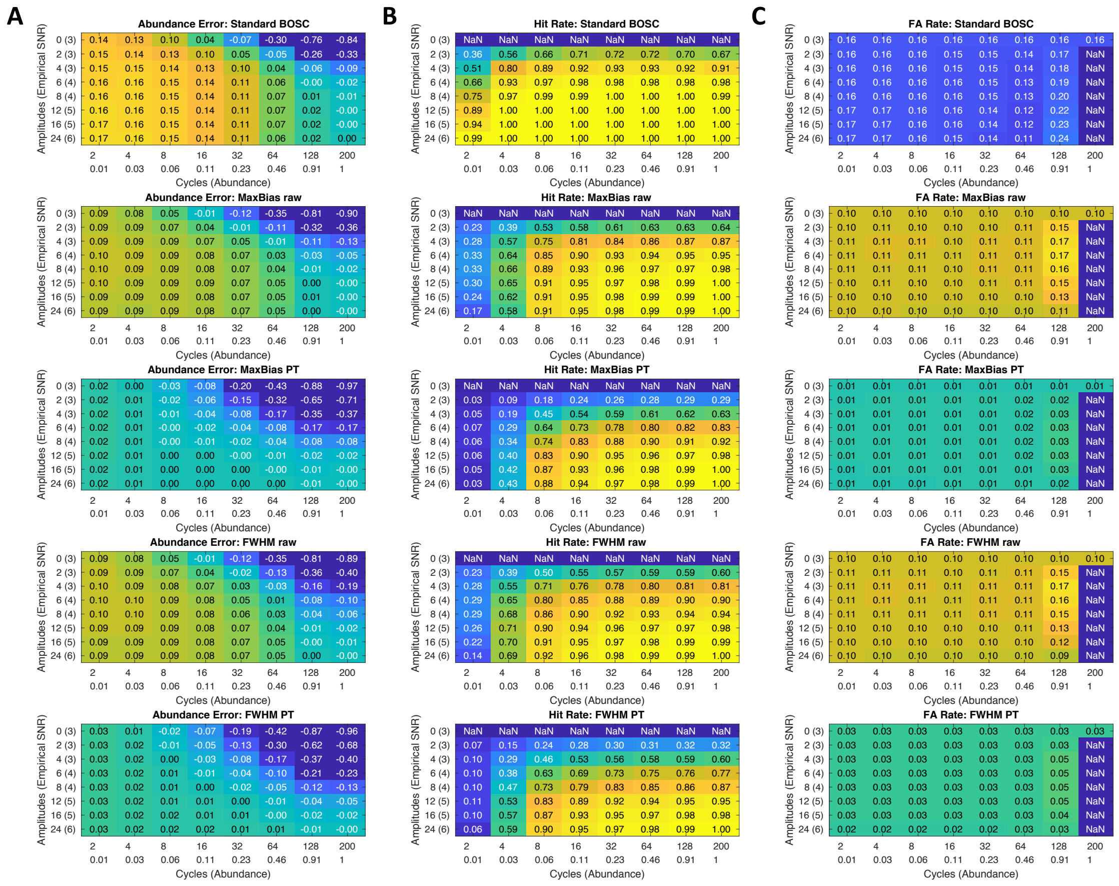

**Fig. S2.** Rhythm detection performance of different post-processing options. (A) Deviation of abundance estimates from simulated duration. (B) Hit rates. (C) False alarm rates. The subplot structure is the same as in Figure 3A. Rows show detection performance for different routines. The analyses in the main paper use the 'MaxBias PT' method. In the 'PT' method, only power values above the threshold were considered for post-processing, otherwise 'raw' power values were considered. Note that using the PT method, abundance is highly specific and is never over- but only underestimated, thus generally providing a lower bound on the real rhythmic abundance.

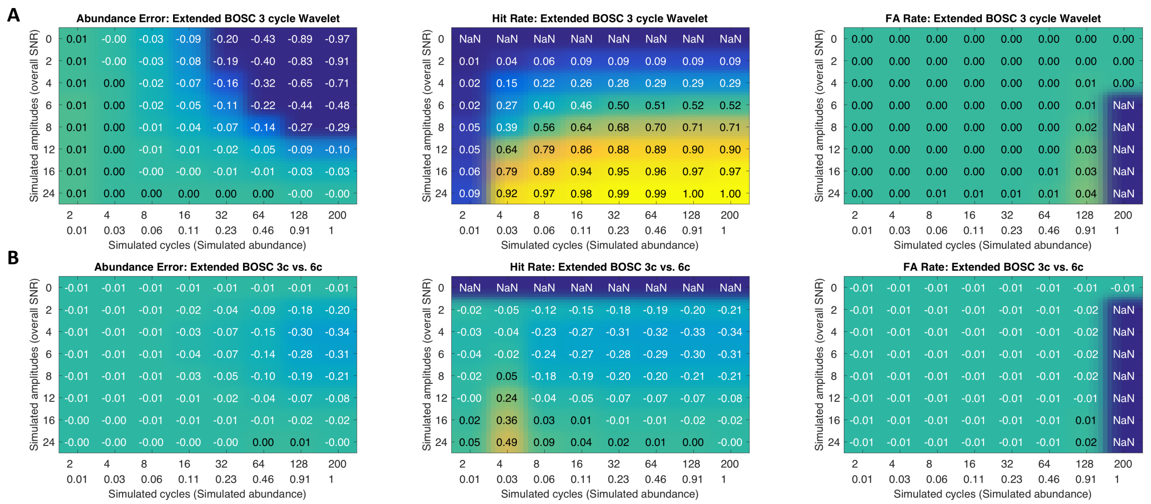

**Fig. S3.** Signal detection results using a 3 cycle wavelet (compared to the 6 cycle wavelet used for the main analyses). (A) Results for data from a 3 cycle wavelet transform indicate high specificity, with a gradient of sensitivity spanned by overall SNR. (B) Compared to the 6 cycle wavelet, the use of a 3 cycle wavelet increases sensitivity for shorter rhythms (around 4 cycles) at high SNR, whereas it decreases sensitivity for more sustained rhythmicity particularly in lower SNR ranges. Specificity is relatively unaffected.

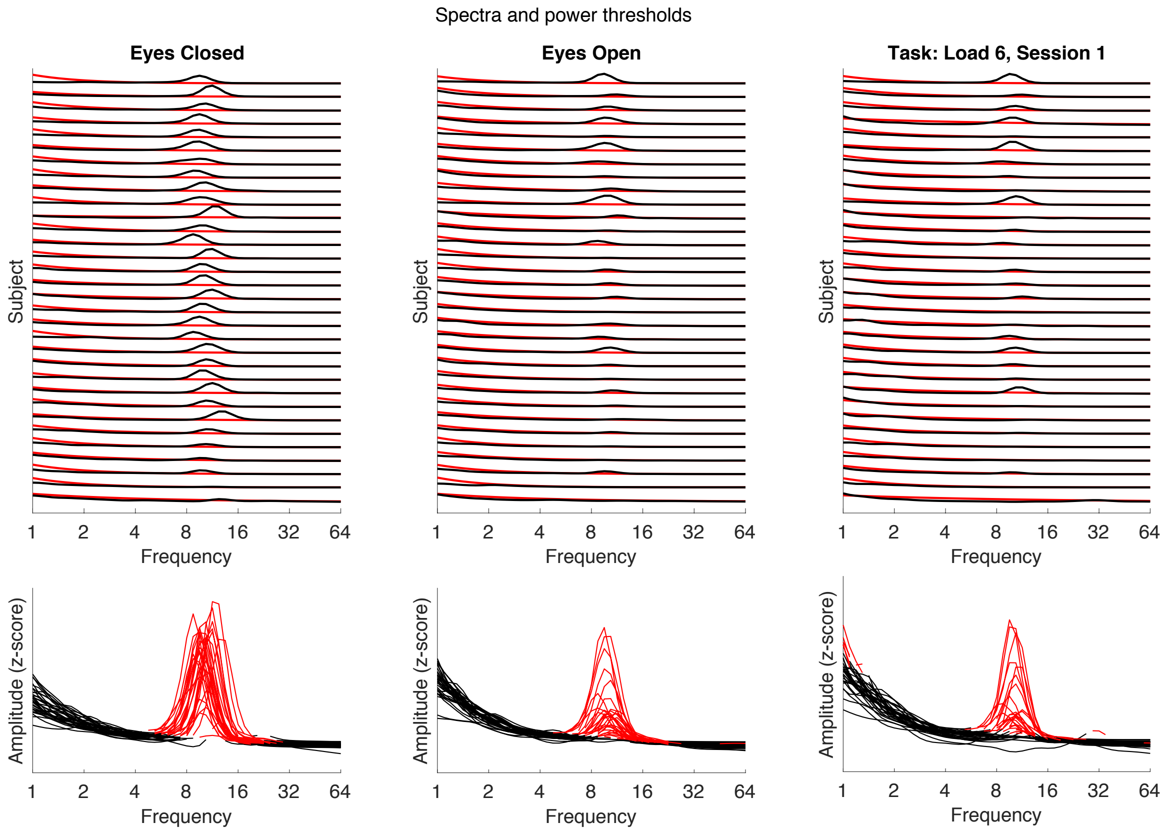


**Fig. S4.** Individual spectra (black) and power thresholds (red) for the eyes closed and eyes open resting state, as well as task retention period. Power thresholds suggest a successful exclusion of the alpha peak and similar fits across subjects. Large spectral peaks are consistently found in the 8-15 Hz range. Spectra are averaged across posterior-occipital channels. Spectra and power thresholds have been spectrally concatenated and z-scored across frequencies for enhanced visibility. Subjects have been sorted by descending 8-15 Hz power during the eyes closed resting state. (Bottom) The alpha peak is consistently excluded from the background threshold. Red traces indicate the spectral power above the power threshold, black traces indicate the segments below the power threshold. Note that falling below the power threshold does not prevent detection as single-trial power can exceed the average power and thus the power threshold. These fluctuations are crucial for the detection of rhythmicity as the power threshold is fixed (see also Figure 7 in main text).

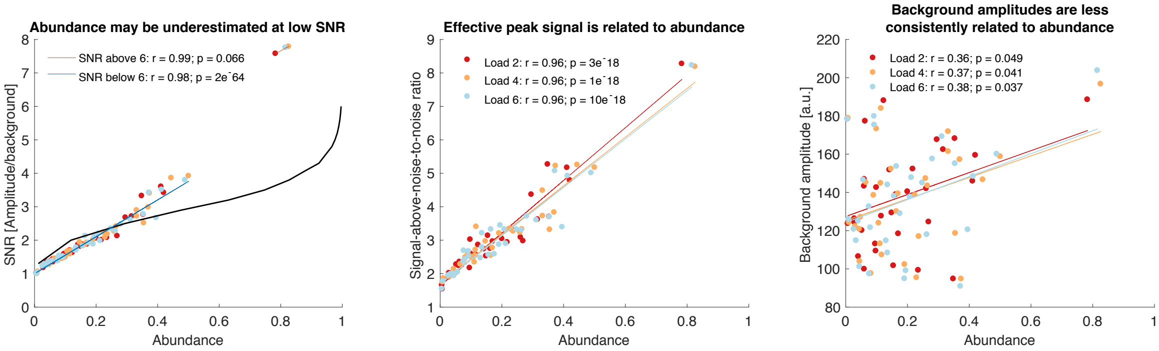


**Fig. S5.** Amplitude-abundance coupling during the retention phase. Similar to the observations in the resting state data (Figure 6 in the main text), the effective rhythmic peak explains the estimated abundance, whereas the background estimate is less consistently associated with abundance

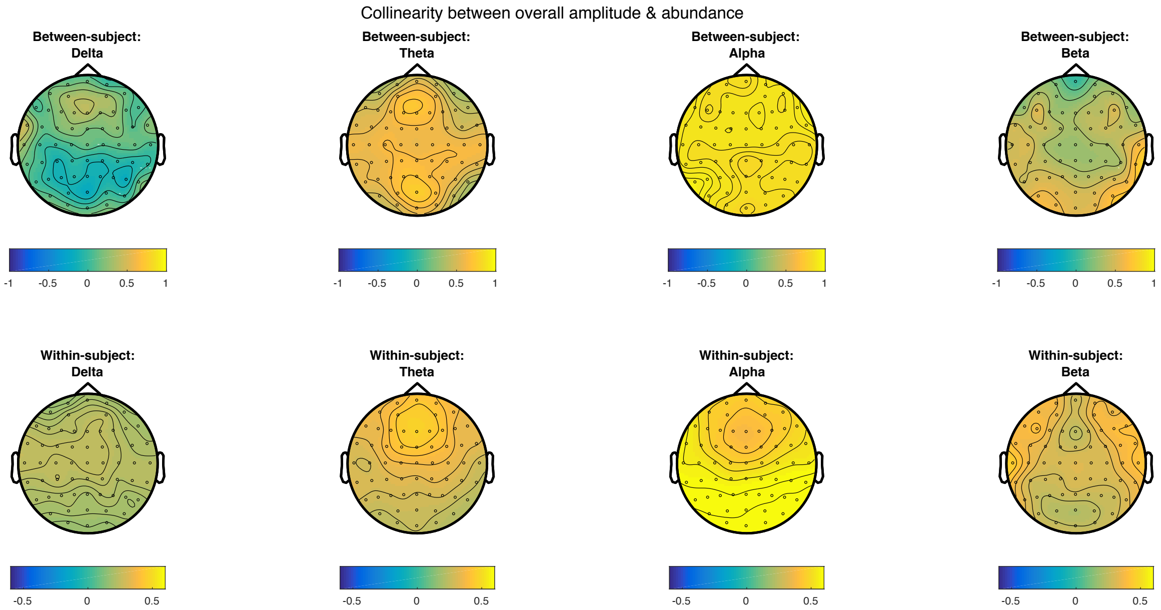


**Fig. S6.** Topographies of between-subject (1st row) and within-subject (2nd row) collinearity (Pearson correlations) of overall amplitude and abundance for multiple low-frequency ranges. Collinearity is highest for the alpha band. Highest within-subject collinearity is observed for channels with high abundance. Corresponding grand average topographies of overall amplitude and abundance are shown in Figure S8.

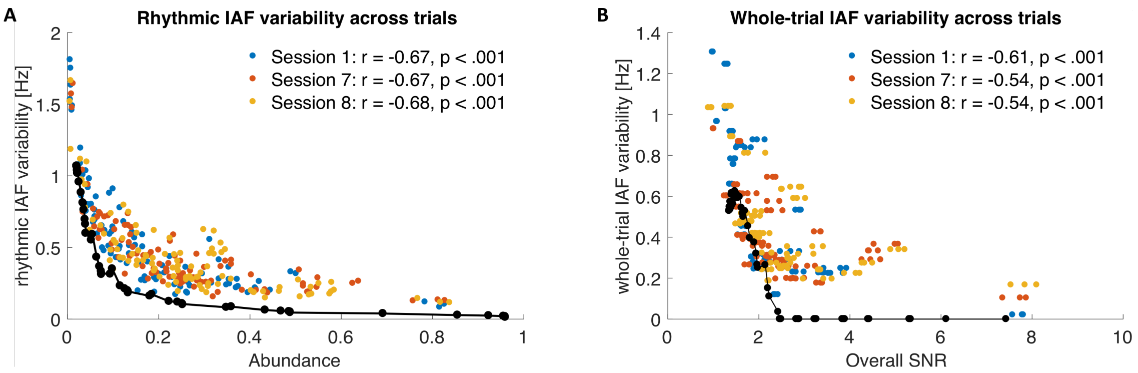


**Fig. S7.** Trial-by-trial IAF variability is associated with sparse rhythmicity. (A) Individual alpha frequency (IAF) precision across trials is related to abundance. Lower individual abundance estimates are associated with increased across-trial IAF variability. (B) This relationship also exists when considering overall SNR and IAF estimates from across the whole trial. Superimposed black lines show the $6^{th}$ order polynomial fit for simulation results encompassing varying rhythm durations and amplitudes. Empirically estimated frequency variability is quantitatively similar to the bias observed at low SNRs in the simulated data.

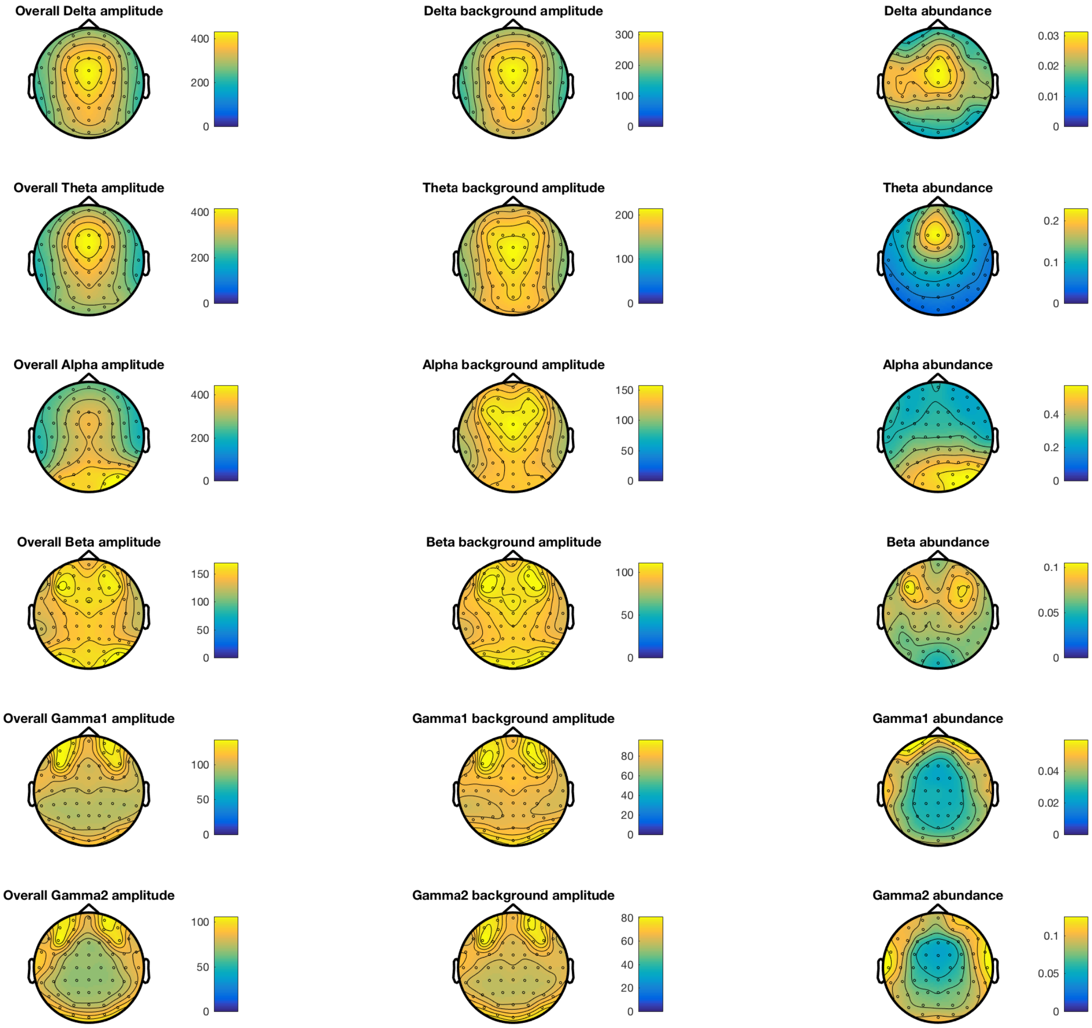


**Fig. S8.** Topographies of overall (narrowband) amplitudes, estimated background amplitudes and abundance indicate a principled exclusion of stationary scale-free background amplitudes from rhythmicity estimates across multiple frequencies. In addition, the topographies suggest that overall amplitudes represent a mixture of a relatively stationary background and spectrally varying rhythmic components. As in Figure 8, the topographies are grand averages from the retention phase of the Sternberg task across sessions.

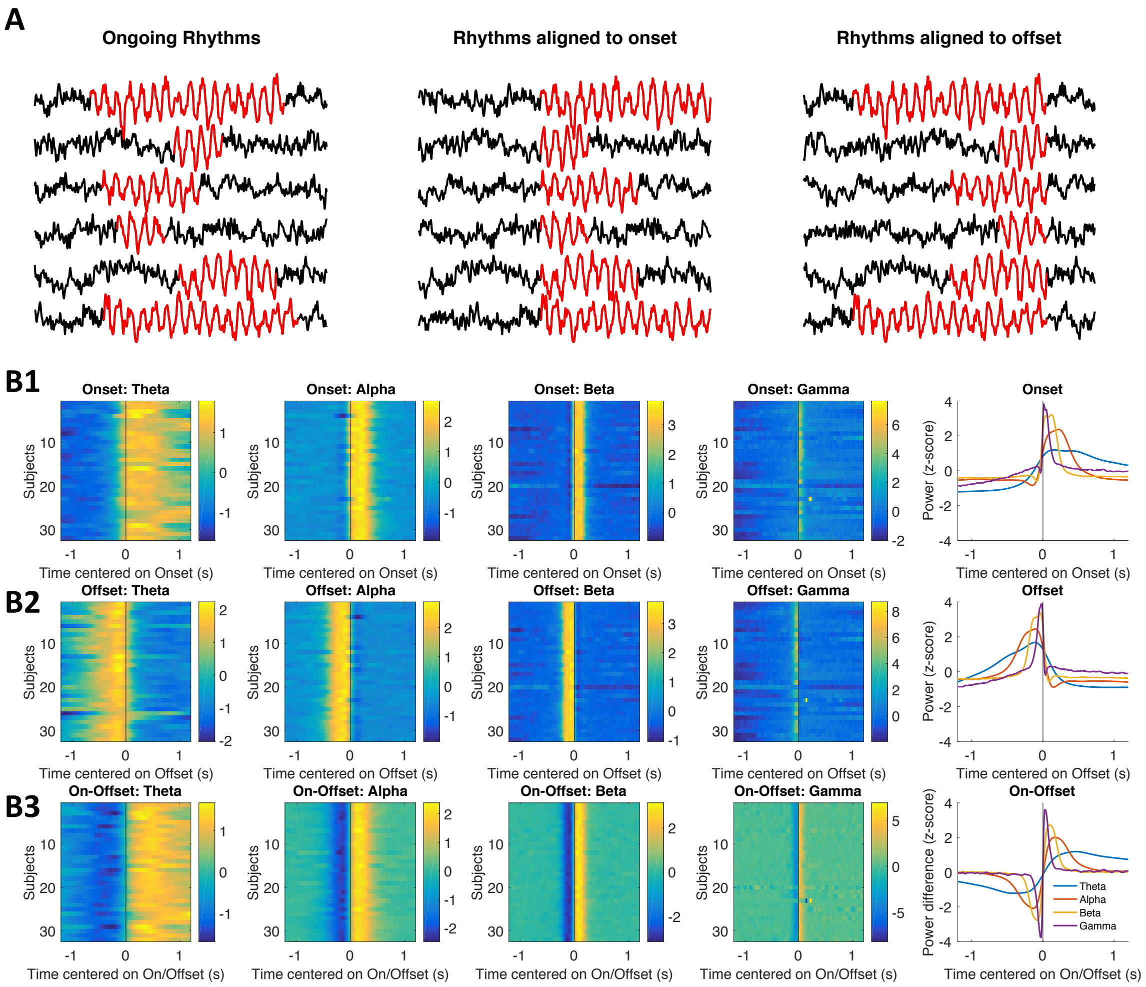


**Fig. S9.** On- and offsets of rhythmic episodes characterize 'rhythm-evoked' effects. (A) Schematic alignment of data to the on- and offsets of rhythmic periods. (B) Rhythm on- and offsets are marked by sudden power shifts at their respective frequency. Individual normalized wavelet power shows a strong increase at the rhythmic onset (B1) and a decrease once rhythmic episodes end (B2). The difference between on- and offset-related power summarizes the evoked effect of rhythmic episodes on ongoing power (B3). Power was extracted within a fixed peri-onset and peri-offset window for all channels where episodes were detected and subsequently averaged across episodes, loads and channels. Finally, the individual averages were z-normalized. The rightmost plots show the grand average across subjects. Data are from extended periods of the Sternberg task in Session 1.

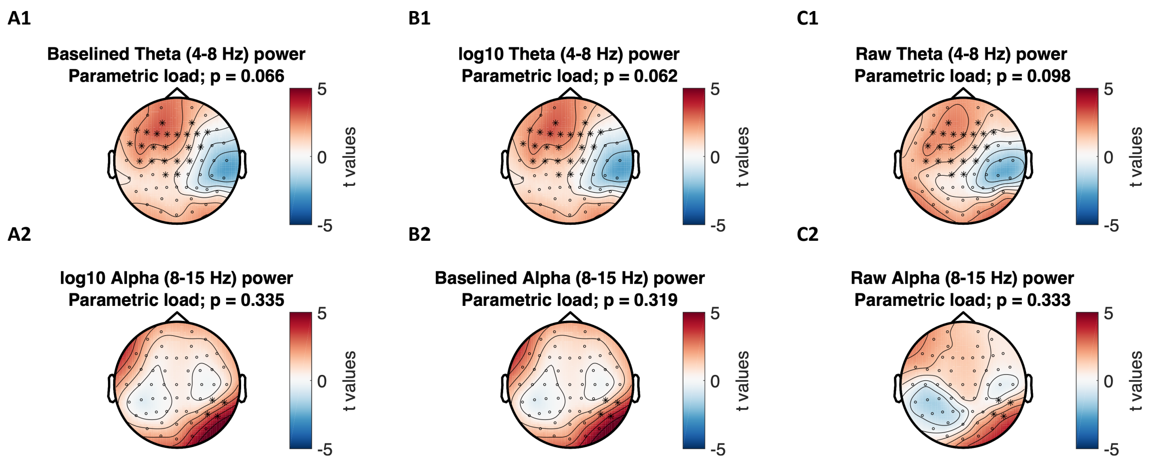


**Fig. S10.** Results of cluster-based permutation tests for different baseline variants: (A) average baseline, (B) single-trial log10-transform, and (C) raw power. While all normalizations produce similar clusters to the observed abundance effect (Figure 11 in main text), no cluster reached statistical significance.

**Table S1.** Berger effect of eye closure on rhythmic and arrhythmic alpha indices, controlling for the high collinearity between indicators

| Dependent variable | Predictor | b | Bootstrap 95% CI Low | Bootstrap 95% CI High | SE | F-value | p-value |
|---|---|---|---|---|---|---|---|
| Rhythmic abundance | Arrhythmic amplitude | 0.33 | 0.15 | 0.51 | 0.09 | 14.0 | 0.001 |
| | Berger effect | −1.11 | −1.44 | −0.79 | 0.16 | 47.7 | 0.000 |
| Rhythmic abundance | Rhythmic amplitude | 0.46 | 0.28 | 0.63 | 0.09 | 26.3 | 0.000 |
| | Berger effect | −0.94 | −1.25 | −0.63 | 0.15 | 36.5 | 0.000 |
| Rhythmic amplitude | Arrhythmic amplitude | 0.85 | 0.76 | 0.95 | 0.05 | 331.4 | 0.000 |
| | Berger effect | −0.26 | −0.42 | −0.10 | 0.08 | 10.4 | 0.003 |
| Rhythmic amplitude | Rhythmic abundance | 0.51 | 0.28 | 0.75 | 0.11 | 19.3 | 0.000 |
| | Berger effect | −0.37 | −0.76 | 0.02 | 0.19 | 3.5 | 0.070 |
| Arrhythmic amplitude | Rhythmic abundance | 0.44 | 0.14 | 0.73 | 0.15 | 8.6 | 0.006 |
| | Berger effect | −0.37 | −0.89 | 0.15 | 0.25 | 2.0 | 0.165 |
| Arrhythmic amplitude | Rhythmic amplitude | 0.99 | 0.88 | 1.09 | 0.05 | 370.3 | 0.000 |
| | Berger effect | 0.10 | −0.09 | 0.29 | 0.09 | 1.0 | 0.323 |

Effects were estimated within linear mixed effects models. Green shading indicates significant ($p < .05$) Berger effects of eye opening that cannot be explained by potential collinearity with the remaining predictor variable; orange shading indicates the absence of an indicated unique Berger effect on the dependent variable.

**Table S2.** Unique memory load effects on rhythm-specific estimates, controlling for high collinearity with traditional estimates

| Dependent variable | Predictor | F-value | p-value |
|---|---|---|---|
| Alpha rhythmic abundance | Alpha overall amplitude | 217.8 | <0.001 |
| | Load | 12.1 | <0.001 |
| Theta rhythmic abundance | Theta overall amplitude | 75.6 | <0.001 |
| | Load | 3.3 | 0.045 |
| Alpha rhythmic abundance | Alpha rhythmic amplitude | 112.4 | 0.000 |
| | Load | 13.5 | 0.000 |
| Theta rhythmic abundance | Theta rhythmic amplitude | 59.8 | 0.000 |
| | Load | 5.3 | 0.008 |
| Alpha rhythmic amplitude | Alpha overall amplitude | 81.0 | <0.001 |
| | Load | 1.3 | 0.278 |
| Theta rhythmic amplitude | Theta overall amplitude | 56.7 | <0.001 |
| | Load | 0.3 | 0.744 |

Effects were estimated within linear mixed effects models. Green shading indicates significant ($p < .05$) memory load effects that cannot be explained by potential collinearity with the remaining predictor variable; orange shading indicates the absence of an indicated unique memory load effect on the dependent variable. Overall amplitudes refer to baselined alpha power as depicted in Figure 11A.
